# B lymphocytes that enter the germinal center late preferentially differentiate into memory cells that recognize subdominant epitopes

**DOI:** 10.1101/2025.10.30.685663

**Authors:** Pengcheng Zhou, Harald Hartweger, Andrew J MacLean, Victor Ramos, Kai-hui Yao, Brianna Hernandez, Zijun Wang, Anna Gazumyan, Michel C. Nussenzweig

## Abstract

Immune responses to pathogens and effective vaccines elicit germinal center (GC) responses wherein B cells undergo affinity maturation and develop into plasma cells (PCs) and memory B cells (MBCs). The GC reaction is initially seeded by a limited group of founder B cells, and subsequently further diversified by continual entry of naïve B cells that compete with GC founder cells for antigen and T cell help. Whether these later-arriving invaders contribute to the development of PCs or MBCs is not known. To investigate the fate of GC invaders we developed a dual-recombinase reporter approach that enables pre- and post-GC B cell lineage tracing and used it to examine immune responses to vaccination and influenza infection. Notably, fate-mapped invaders preferentially give rise to MBCs as opposed to PCs. Moreover, antibodies expressed by invader-derived MBCs harbor fewer somatic mutations, exhibit lower affinity, and their antibodies bind to subdominant antigenic epitopes relative to founder MBCs. Our findings indicate that invader GC B cells are an important source of humoral immune memory diversification after infection or vaccination.

## Introduction

Humoral immune responses are characterized by a rapid increase in the affinity of antibodies in plasma. Affinity maturation is mediated by B lymphocyte clonal expansion, somatic mutation and selection in GCs.^1,2^ These microanatomical structures produce PCs that secrete the antibodies found in circulation and MBCs that can be called upon when the host is re-challenged with the immunogen or pathogen. In parallel to increased affinity, another prototypical feature of the antibody response is increasing diversity, a phenomenon first noted by Landsteiner in 1936, and underscored by Eisen in studies on immune responses to haptens.^3,4^

Rapid diversification is essential for mounting effective antibody responses against evolving pathogens and their variants such as influenza, and SARS-CoV-2.^5-8^ Diversification occurs at several different points in the response beginning with recruitment of a large number of different B cells expressing antibodies of varying affinity into GCs.^9-11^ This initial group of founder GC B cells receive positive selection signals from antigen and specialized T follicular helper cells (T_FH_) and undergo clonal expansion and proliferative bursting which focuses the response at the cost of reduced clonal diversity.^10,12^ Recent studies show that GC cellular diversity is maintained in part by later arriving naïve invader B cells that generally show lower initial levels of antibody affinity than founder cells.^13-15^ How these invaders might contribute to PC and MBC compartments is not known.

Here, we report on dual reporter lineage-tracing experiments that fate-map the development of founder and invader GC B cells into PC and MBCs. The data indicate that naïve B cells entering the GC in the late stages of the response diversify the immune response by preferentially developing into MBCs that express antibodies targeting subdominant epitopes.

## Results

### Dynamics of PC and MBC production

To investigate the dynamics of MBC and PC development in a polyclonal immune response irrespective of rates of post GC cell division, we immunized *S1pr2*^CreERT2/+^ *Rosa26*^LSL-zsGreen/+^ mice ^16^ with 4-hydroxy-3-nitrophenylacetyl hapten conjugated to ovalbumin (NP-OVA) (**Figure 1A**). The progeny of activated B cells were permanently labeled by weekly administration of tamoxifen starting on day 5, at the beginning of the GC reaction and after termination of the initial burst of extrafollicular MBC and PC development.^17^ Tamoxifen induces deletion of the STOP cassette in the *ROSA* locus and permanently labels GC B cells and their progeny with zsGreen (zsG) (**Figure 1A**). Popliteal lymph node (LN) GC, MBC, and PC populations were analyzed on days 11, 18, 25, and 32 (**Figure 1A-C and Figure S1A**). As expected, 90-95% of all GC B cells were labeled at all time points assayed (**Figure 1B**). Overall GC B cell and PC numbers declined rapidly from a peak at day 11 to day 32 (**Figure 1C**). By contrast, the number of labeled MBCs (zsG^+^CD38^+^Fas^-^) in the draining lymph node was stable over this period (**Figure 1C**).

**Figure 1.**
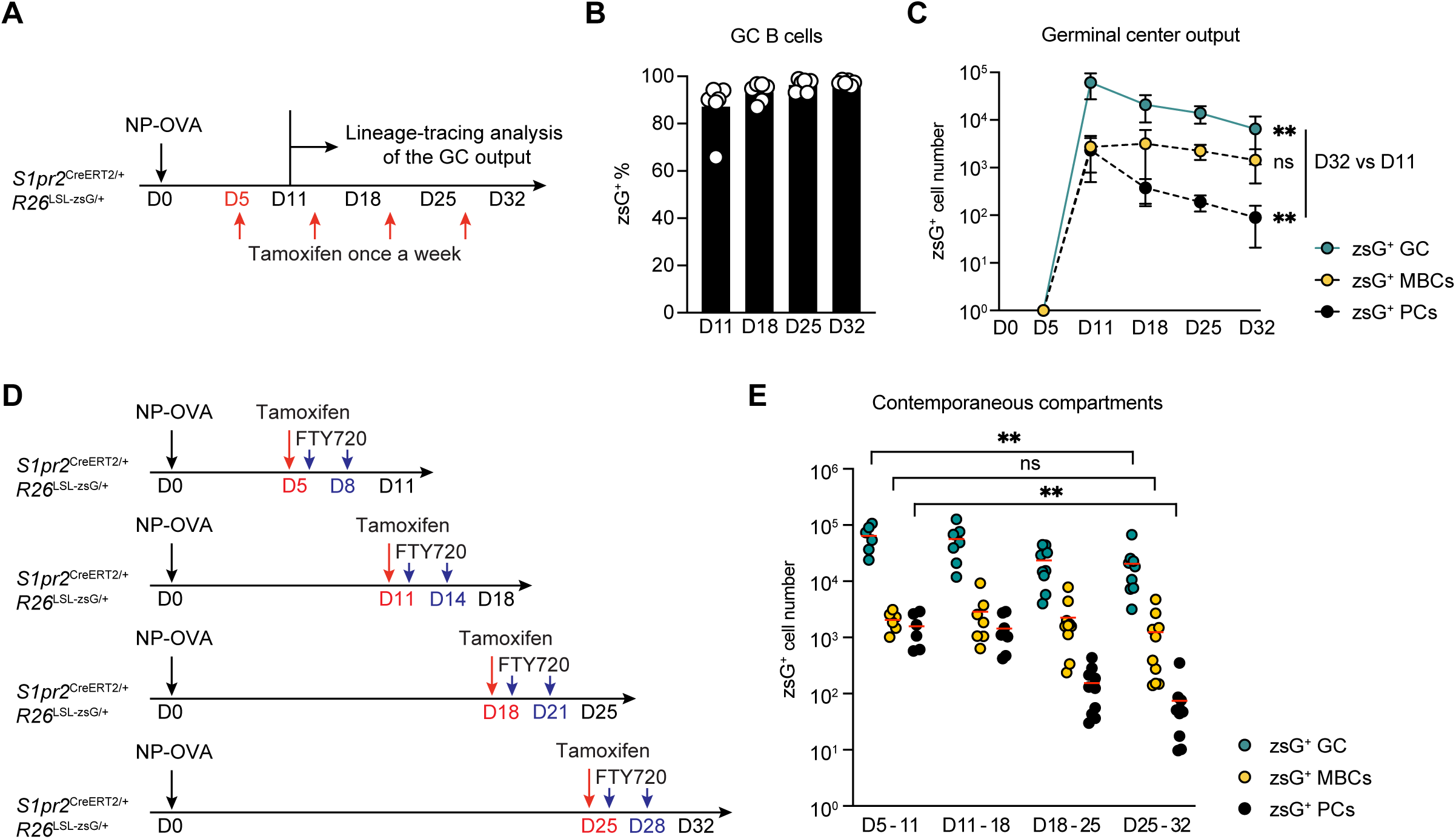
Temporal dynamics of germinal center output. **(A)** Schematics of the experiment. *S1pr2*^CreERT2/+^ *Rosa26*^LSL-zsGreen/+^ mice were immunized with NP-OVA in footpad on day 0, treated with tamoxifen on days 5, 12, 19, and 26. Popliteal lymph nodes were obtained, and B220^+^CD38^-^Fas^+^ GC B cells, B220^+^CD38^+^Fas^-^zsG^+^ MBCs and Dump^-^ TACI^+^CD138^+^ plasma cells were analyzed on days 11, 18, 25, and 32. **(B)** Frequency of zsGreen^+^ GC B cells labelled by tamoxifen administration at each time points. Bar indicates mean. **(C)** Graph showing the cell numbers of zsGreen^+^ GC B cells (blue), MBCs (yellow) and plasma cells (black) over time. Circle indicates mean, and error bars indicate SD, n = 6 at each time point. P values calculated using one-way ANOVA, *P ≤ 0.05, **P ≤ 0.01, two independent experiments. **(D)** Pulse-chase experiment setup. *S1pr2*^CreERT2/+^ *Rosa26*^LSL-zsGreen/+^ mice were immunized with NP-OVA in footpad on day 0, treated with tamoxifen on days 5 or 11, or 18 or 25 to label the GC B cells in each window. Cell egress from draining lymph node was prevent by two times FTY720 injection in each window, and GC B cells, MBCs and plasma cells were analyzed on days 11, 18, 25, and 32. **(E)** Graph showing the cell numbers of zsGreen^+^ GC B cells (blue), MBCs (yellow) and plasma cells (black) on days 11, 18, 25, and 32 after NP-OVA immunization. Red line indicates mean. Statistics on zsGreen^+^ GC B cells (blue), MBCs (yellow) and plasma cells (black) between day 5-11 to day 25-32. P values calculated using one-way ANOVA, *P ≤ 0.05, **P ≤ 0.01, three independent experiments.

The observed differences might reflect preferential differentiation of GC B cells into MBCs or PCs over time, or differences in lymph node retention versus egress between MBCs and PCs. To determine whether PC and MBC production changes during the immune response, we performed pulse-chase experiments. Lymph node egress by newly developing PCs and MBCs was inhibited using FTY720 to block S1PR1 (**Figure 1D-E** and **S1B**).^18^ A single pulse of tamoxifen was administered on days 5, 11, 18, or 25 after immunization, and labeled GC B cells, MBCs and PCs were enumerated one week later. Whereas the number of labeled GC B cells decreased over time, the number of MBC produced remained relatively stable (**Figure 1E**). In contrast, contemporaneously labeled PCs waned rapidly, exhibiting a mean of 21-fold reduction in the day 25-32 window compared with the day 5-11 (p= 0.001, **Figure 1E**). Thus, production of PCs peaks early and decreases rapidly while MBC production increases relative to the size of the GC during the response.

### Dual reporter lineage-tracing system to track GC invaders

Tamoxifen administration to *Cd55*^CreERT2/+^ *Rosa26*^LSL-tdTomato/+^ reporter mice labels naïve follicular but not GC B cells or PCs enabling fate mapping of naïve B cells entering the GC during later stages of the reaction (**Figure S2A**).^13,19^ However, in addition to naïve B cells, MBCs also express high-levels of CD55 which makes it impossible to determine how naïve B cells entering the GC in the late stages of the reaction contribute to the memory compartment over time (**Figure S2B**).^19^

To address this shortcoming and examine the contribution of B cells recruited at later stages of the immune response ^13-15,20^ to the PC and MBC compartment, we developed a dual-recombinase reporter system. Mice harboring a novel *Cd55*^L-DreERT2-hCD2-L/+^ *Rosa26*^RSR-tdTomato/+^ reporter were combined with *Aicda*^Cre/+^ *Rosa26*^LSL-zsGreen/+^ mice ^21^ (CD55^DreER^AID^Cre^ dual reporter mice, **Figure 2A and Figure S2C**). In these mice, human CD2 (hCD2) reports the presence of tamoxifen-inducible Dre recombinase (Dre^ERT2^) in the *Cd55* locus **(Figure S2D-E)**. The Dre^ERT2^-hCD2 cassette is additionally flanked by loxP sites for Cre recombinase-mediated deletion. Tamoxifen administration removes the Rox-flanked STOP cassette from the *Rosa26*^RSR-tdTomato^ allele allowing tdTomato (tdT) expression in naïve B cells. Constitutive Cre recombinase expression under the control of *Aicda* labels GC B cells with zsGreen by deleting loxP-flanked STOP cassette from the *Rosa26*^LSL-zsGreen/+^ locus and removes the loxP-flanked Dre^ERT2^-hCD2 sequence from the *Cd55* locus (**Figure 2A**).

**Figure 2.**
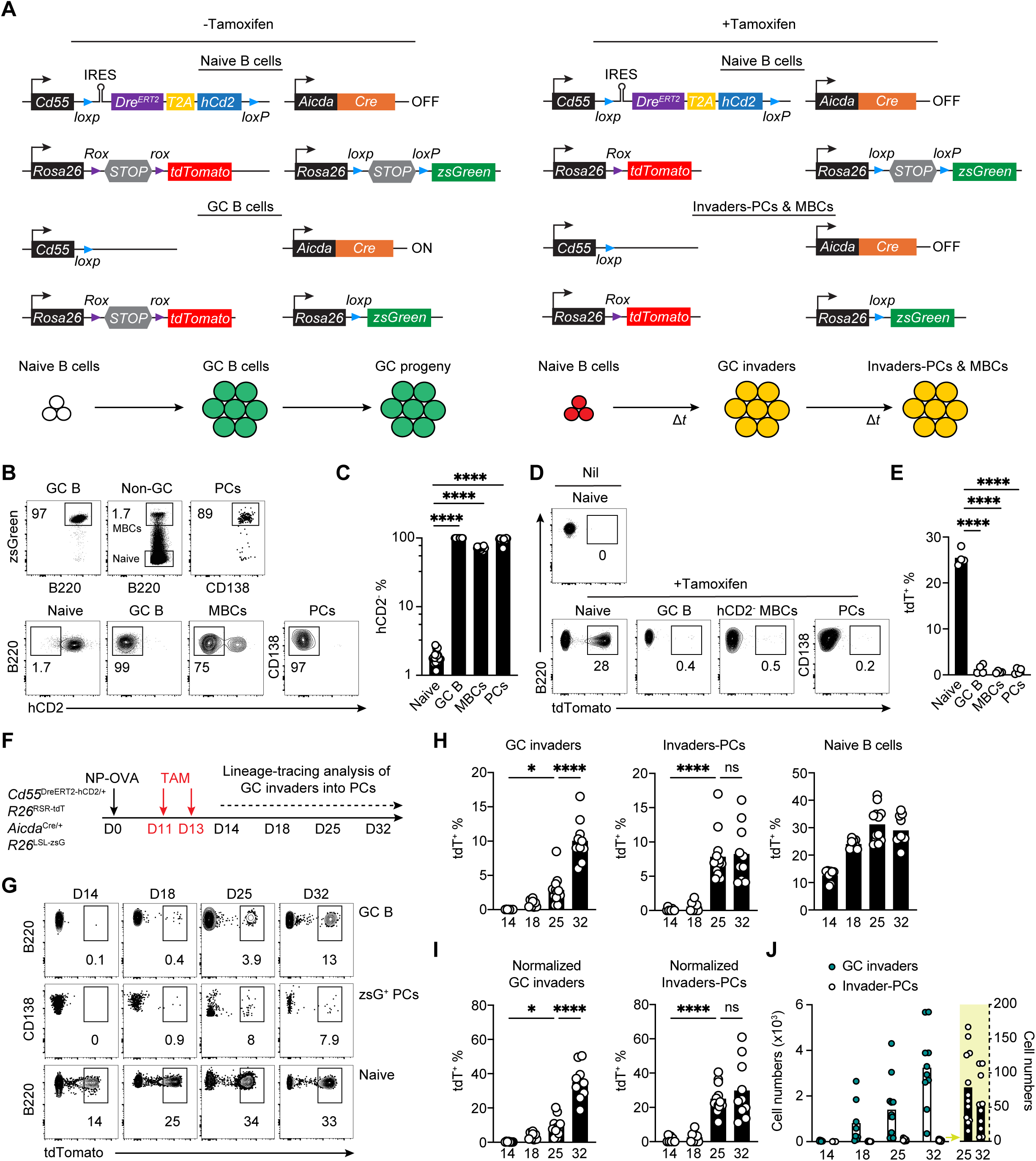
Lineage tracing system of invader B cells and fate-mapping into PC compartment. **(A)** Schematics of the CD55^DreER^AID^Cre^ dual reporter mouse model. Top, genomic architecture of reporter alleles. Bottom, depiction of fluorescent protein expression in cells before and after tamoxifen administration. **(B)** Representative flow cytometry plots showing the zsGreen expression in LN B220^+^CD38^-^Fas^+^ GC B cells, non-GC B cells and Dump^-^TACI^+^CD138^+^ PCs, and hCD2 expression in naïve, GCs, MBCs and PCs isolated from CD55^DreER^AID^Cre^ dual reporter mice with NP-OVA immunization. **(C)** Graph showing the percentage of hCD2^-^ cells in naïve, GCs, MBCs and PCs. Each dot represents one mouse. Bar indicates mean. **(D)** Representative flow cytometry plots showing the tdTomato expression in control naïve B cells or naïve B cells, GC B cells, MBCs and PCs 7 days after treatment with tamoxifen; MBCs were gated on hCD2^-^zsG^+^, and PCs were zsG^+^. **(E)** Statistics of tdTomato^+^ cells in **(D)**. Each dot represents one mouse, bar indicates mean, one-way ANOVA, ****P ≤ 0.0001. **(F)** Schematics of the experiment. CD55^DreER^AID^Cre^ dual reporter mice were immunized with NP-OVA in footpad on day 0, and treated with tamoxifen on day11 and 13. Popliteal lymph nodes were obtained, and GCs and PCs were analyzed on days 14, 18, 25, and 32. **(G)** Representative flow cytometry plots summarizing the percentage of tdTomato^+^ cells in naïve, GC and zsG^+^ PCs on days 14, 18, 25 and 32. **(H)** Statistics of tdTomato^+^ cells in naïve, GC and PCs show in (**G**). **(I)** Statistics of tdTomato^+^ cells in GC and zsG^+^ PCs after correcting for labeling efficiency of the naïve B cell compartment. **(J)** Number of tdTomato^+^ cells in GCs (Green, GC invaders) and PCs (White, Invaders-PCs). Yellow area highlighting the PC counts on a different scale. At least 2 independent experiments, each dot represents one mouse, bar indicates mean, n=8-12, one-way ANOVA, ****P ≤ 0.0001.

In the absence of tamoxifen, B cells expressing *Aicda* and their PC and MBC progeny should express zsGreen and delete the Dre^ERT2^-hCD2 in the *Cd55*^L-DreERT2-hCD2-L/+^ locus (tdT^-^hCD2^-^zsG^+^). Consequently, cells that have expressed *Aicda* lose the ability to express Dre^ERT2^ and cannot respond to tamoxifen administration by removing the Rox-flanked STOP cassette from the *Rosa26*^RSR-tdTomato/+^ allele to induce tdTomato expression and they instead remain tdT^-^hCD2^-^zsG^+^. In contrast, naïve B cells would be labeled with tdTomato after tamoxifen administration, and upon entering GCs these cells would also delete the STOP cassette in the *Rosa26*^LSL-zsGreen/+^ locus and co-express zsGreen (tdT^+^hCD2^-^zsG^+^) (**Figure 2A**).

To test the dual recombinase system, we immunized dual-reporter mice with NP-OVA and assayed draining lymph nodes by flow cytometry. Fourteen days after immunization, 99% of the GC B cells, 60-80% of all MBCs and over 95% of PCs in draining LNs were tdT^-^hCD2^-^zsG^+^, indicating high-efficiency of *Aicda*^Cre/+^ mediated excision of both the Dre^ERT2^-hCD2 from the *Cd55* locus and the STOP cassette from the *Rosa26*^LSL-zsGreen^ locus (**Figure 2B and C**). Upon tamoxifen injection, naïve dual reporter B cells should excise the STOP cassette from the *Rosa26*^RSR-tdTomato/+^ locus and become tdT^+^hCD2^+^zsG^-^ (**Figure 2A**). Seven days after tamoxifen administration, 25-30% of naïve B cells were tdT^+^hCD2^+^zsG^-^, whereas tdT^+^hCD2^-^zsG^+^ MBCs and PCs were barely detectable at this time (**Figure 2D and E**). tdT^+^hCD2^+^zsG^-^ naive B cells that enter the GC after tamoxifen administration will remain tdT^+^, express *Aicda*^Cre^, and should become zsG^+^ (**Figure 2A**). They would also be expected to excise the lox-flanked Dre^ERT2^-hCD2 sequence from the *Cd55* locus and become tdT^+^hCD2^-^zsG^+^. hCD2⁺zsG⁺ cells were always excluded to avoid any confounding tdT⁺ labeling of pre-existing MBCs. In this way, zsG expression labels all progeny of activated B cells and, separately, tdT expression reports all progeny of cells that were naïve at the time of tamoxifen administration. Thus, this system enables us to distinguish between tdT^-^hCD2^-^zsG^+^ founder and tdT^+^hCD2^-^zsG^+^ invader origin PCs or MBCs respectively (**Figure 2A**).

### PCs developing from B cells entering late

To determine whether naïve B cells that become activated in the later stages of the immune response develop into PCs, tamoxifen was administered to dual reporter mice 11 and 13 days after immunization with NP-OVA (**Figure 2F, Figure S2F-H, and Figure S3A-D**). Naïve B cell labeling was detected on day 14, 3 days after tamoxifen administration, and in GCs on day 18 (**Figure 2G and H**). Over the following weeks, invader origin tdT^+^hCD2^-^zsG^+^ cells accumulated in the GC and PC compartments, reaching 35.1% and 29.8% of these compartments respectively on day 32 after correction for labelling efficiency of the naïve B cell compartment (**Figure 2H-I**). However, the absolute number of PCs developing from invader GC B cells recruited into the late stages of the reaction was very small (**Figure 2J**). Similar results were obtained after administration of the Tenivac vaccine which is a combination of tetanus and diphtheria toxoid antigens (**Figure S3E**-**G**).

To assess the potential extrafollicular response that may be marked by the dual reporter system, CD55^DreER^AID^Cre^ mice were immunized with NP-OVA, administered tamoxifen on day 11, and analyzed by confocal microscopy on day 15 (**Figure S4A**). 15 days post immunization, confocal imaging revealed very few tdT⁺zsG⁺ cells in extrafollicular regions of the lymph node (**Figure S4B & 4D**). Among the few tdT⁺zsG⁺ cells, the majority localized around the GC borders and within B cell follicles (**Figure S4B**). Additional dual reporter mice were immunized with NP-OVA and treated with tamoxifen on day 0 and imaged on day 4 at the peak of the extrafollicular response.^17,22^ Consistent with early extrafollicular activation that precedes GC entry, imaging on day 4 post-immunization showed more abundant tdT⁺zsG⁺ cells in extrafollicular areas (**Figure S4C & 4D**). Together, these imaging results confirm that extrafollicular responses are more prevalent in earlier stages of the response.

To characterize the antibodies produced by PCs and contemporaneous GC B cells derived from invader B cells, we isolated individual fate-mapped tdT^+^hCD2^-^zsG^+^ invader-origin and tdT^-^hCD2^-^ zsG^+^ founder cells from contemporaneous GCs and PCs. CD55^DreER^AID^Cre^ indicator mice were immunized with NP-OVA, tamoxifen was administered on days 11 and 13 and cells were isolated on day 25 after immunization (**Figure 2F**). At this time tdT^-^hCD2^-^zsG^+^ cells represent a mixture of approximately 90% of bona-fide founders and 10% invader cells that failed to delete the Rox-floxed STOP cassette in the *ROSA* locus. (**Figure S5A-B)**. PCs derived from B cells entering the later stages of the response (tdT^+^hCD2^-^zsG^+^) were more diverse, less clonal and carried fewer somatic mutations than controls (**Figure 3A-E and S5C**). As an initial surrogate for affinity, we enumerated IGHV1-72 expression which is associated with high-affinity NP binding.^23,24^ On day 25 when invaders comprise 9.1% of all GC B cells after correction for labeling efficiency, 31% and 65% of the control tdT^-^hCD2^-^zsG^+^ GC B cells and PCs respectively expressed IGHV1-72. By contrast, only 7.5% and 5.1% of invader derived tdT^+^hCD2^-^zsG^+^ GC B cells and PCs expressed IGHV1-72 suggesting that invader PCs are less enriched in high affinity clones (p<0.0001, **Figure 3F**).

**Figure 3.**
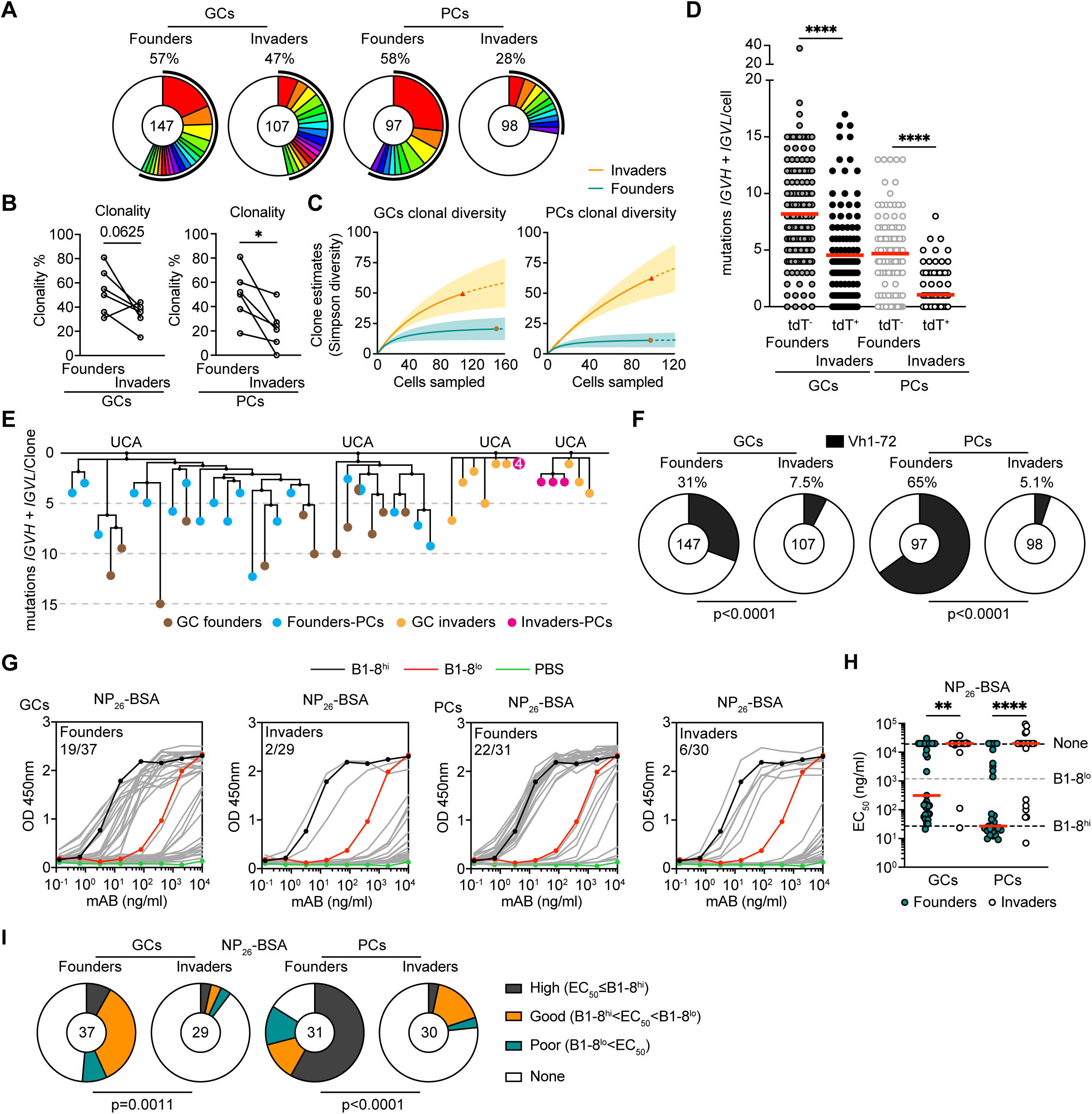
Antibodies produced by invader-derived PCs. **(A)** Pie charts depicting the distribution of antibody sequences (paired VH+VL) for GC founders, GC invaders, Founders-PCs and Invaders-PCs isolated 25 days post NP-OVA immunization of CD55^DreER^AID^Cre^ dual reporter mice. Data pooled from 6 individual mice. The number in the inner circle indicates the number of sequences analyzed. The pie slice size is proportional to the number of clonally related sequences. The black outline and associated numbers indicate the percentage of clonal sequences detected at each time point. **(B)** Graph shows clonality of founders and invaders in GCs and PCs from each mouse. **(C)** Simpson diversity quantifications. (**D**) Number of somatic mutations (nucleotides) of paired VH+VL chains. Each dot represents one cell with paired Ig genes. Red line indicates mean. One-way ANOVA, ****P ≤ 0.0001. (**E**) Representative trees showing phylogenetic relationships between paired VH+VL sequences obtained from GC founders (Brown), Founders-PCs (Blue), GC invaders (Yellow) and Invaders-PCs (Red) identified in **(D)**. The top line represents the clone’s unmutated common ancestor (UCA). **(F)** Pie charts depicting the enrichment of IGHV1-72 clones from Ig gene sequencing analysis of GC founders, GC invaders, Founders-PCs and Invaders-PCs. The pie slice size is proportional to the number of IGHV1-72 clones. The associated numbers indicate the percentage of IGHV1-72 clones. The number in the inner circle indicates the number of sequences analyzed. Two-tailed Fisher’s exact test. **(G)** ELISA binding of 127 monoclonal antibody Fabs derived from each group in **(A)** to NP_26_-BSA. Each grey curve represents one Fab. Left, Fabs produced from GC founders and GC invaders. Right, Fabs produced from Founders-PCs and Invaders-PCs. Control antibodies: high-affinity B1-8^hi^ is in black, low-affinity B1-8^lo^ is in red, PBS control is in green. Numbers show Fabs with EC_50_ below B1-8^lo^ among total in each group. **(H)** Graph shows EC_50_ of Fabs binding to NP_26_-BSA. Each dot represents one Fab. Red line indicates median. One-way ANOVA, **P ≤ 0.01. ****P ≤ 0.0001. **(I)** Pie charts depicting the percentage of high (black)-, good (yellow)-, poor-binding (green) Fabs and non-binders (white) in each GC and PC compartments. The number in the inner circle indicates the number of Fabs analyzed. Two-tailed Fisher’s exact test.

To assess the relative binding properties of the antibodies derived from invader PCs, we produced 127 monoclonal antibody fragment antigen-binding region (Fabs) and tested them for binding to NP-26 bovine serum albumin (BSA) by enzyme-linked immunosorbent assays (ELISA) (**Figure 3G-I**).^25,26^ Whereas 51% and 71% of the Fabs obtained from founder GC B and PCs showed binding activity equal to or greater than the control antibody (B1-8^lo^, *K*_a_ =1.25×10^5^ M^-1^),^23^ only 6.8% and 20.0% of the Fabs obtained from invader GC B and PCs did so respectively (**Figure 3G-I**). When non-binders were excluded from the EC_50_ analysis, Fabs cloned from founder B cells exhibited significantly lower EC_50_ than invaders (**Figure S5D**). In addition, IGHV1-72 usage was significantly enriched among founder-derived Fabs compared to invaders (**Figure S5E**). In summary, naïve B cells entering the late stages of the immune response develop into small numbers of PCs that produce antibodies with significantly decreased ability to bind to the original antigen compared to founders (**Figure 3H and I**).

### Invader-derived memory

To determine whether GC invaders differentiate into MBCs, we immunized dual reporter mice with NP-OVA or a vaccine antigen, SARS-CoV-2 Wuhan-Hu-1 (WT) receptor binding domain (RBD) (**Figure 4A and B, Figure S6A and B**). Mice received tamoxifen on days 11 and 13 to label naïve B cells with tdT (**Figure 4A**). As expected, germinal center size peaked on day 14 and gradually decreased over the next 2.5 weeks (**Figure 4B**). tdT^+^ cells accumulated in the ongoing GCs over the subsequent observation period (**Figure 4C**), reaching a peak of 30%-33% of the GC after correction for labelling efficiency of the naïve B cell compartment (**Figure 4D and E**). GC invaders differentiated into tdT^+^hCD2^-^zsG^+^ MBCs overtime (**Figure 4C**), reaching 20%-25% of the cells in this compartment after correction for labelling efficiency on day 32 after immunization (**Figure 4D and E**). Notably, when comparing total cell numbers, we observed approximately a 10-fold higher number of tdT⁺ cells among hCD2^-^zsG^+^ MBCs than among zsG^+^ PCs at day 25, with this difference further increasing by day 32 (**Figure S6C)**.

**Figure 4.**
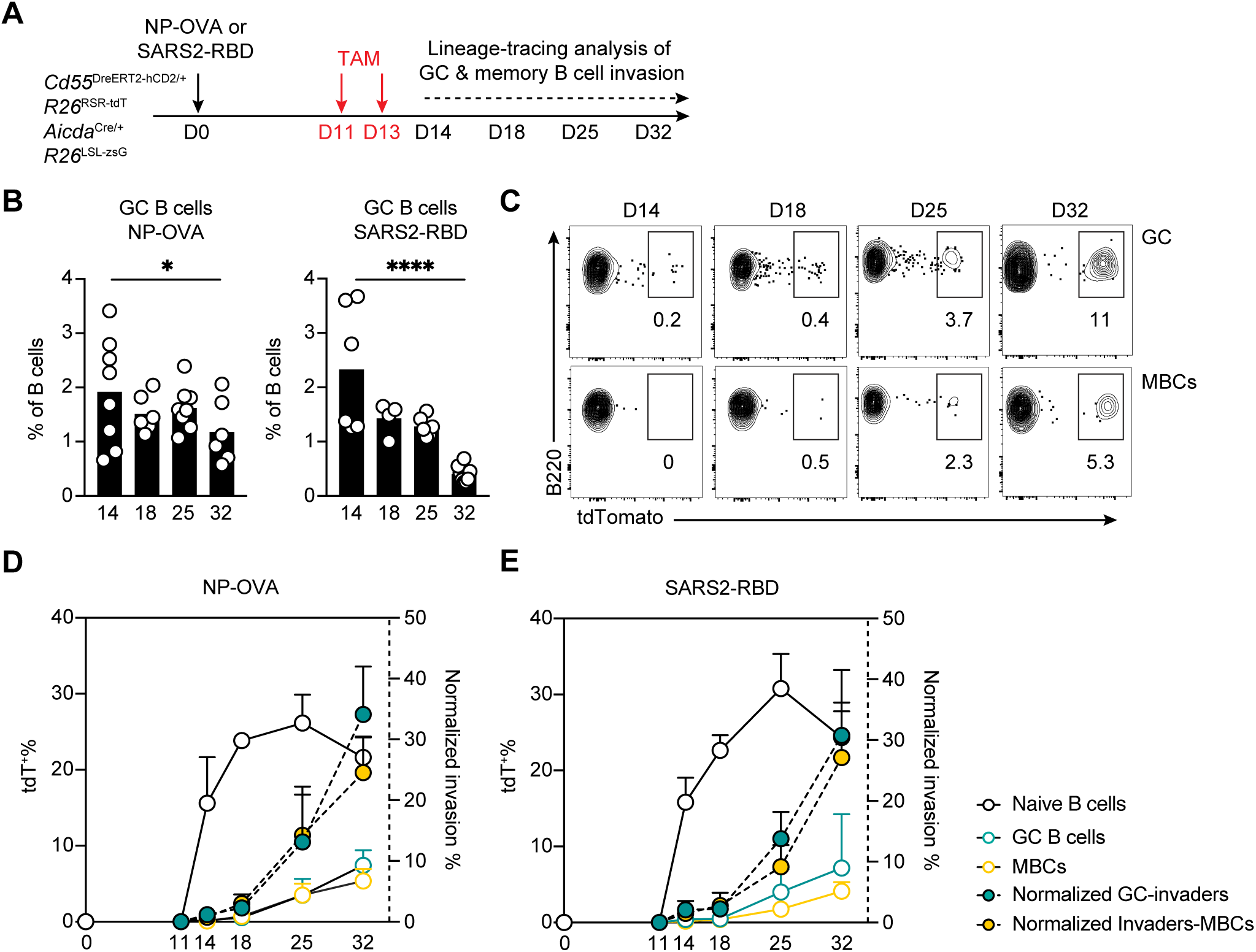
Fate-mapping into memory B cell compartment. **(A)** Schematics of the experiment. CD55^DreER^AID^Cre^ dual reporter mice were immunized with NP-OVA or SARS-CoV-2 RBD protein, administrated with tamoxifen on day 11 and 13 and popliteal lymph nodes (pLN) were assayed on days 14, 18, 25 and 32. **(B)** Graph showing the percentage of GC B cells over time. Each dot represents one mouse, bar indicates mean, 2 or 3 independent experiments, n=4-8 for each time point. **(C)** Representative flow cytometry plots summarizing the percentage of tdT^+^ GCs and tdT^+^hCD2^-^zsG^+^ MBCs on days 14, 18, 25 and 32 in mice immunized with SARS-CoV-2 RBD protein. **(D-E)** Graph showing the percentage of tdT^+^ naïve B cells (white circle), tdT^+^ GC B cells (green circle) and tdT^+^hCD2^-^zsG^+^ MBCs (yellow circle) over time. Green filled dots with dash line show frequency of tdT^+^ GC B cells and yellow filled dots with dash line show frequency of tdT^+^hCD2^-^zsG^+^ MBCs normalized for the labeling efficiency of the naïve B cell compartment. 2 or 3 independent experiments, circle/dot indicates mean, and error bars indicate SD.

To examine the antibodies produced by invader MBCs, we purified tdT^+^hCD2^-^zsG^+^ invader and tdT^-^hCD2^-^zsG^+^ founder MBCs on days 25 and 32 after immunization with NP-OVA and sequenced their antibodies (**Figure 5A**). As expected, both types of MBCs showed relatively low levels of clonality (**Figure 5B**).^27^ However, invader derived memory differed from tdT^-^hCD2^-^zsG^+^ MBCs in showing fewer somatic mutations (day 25, p=0.031; day 32, p=0.001, **Figure 5C-D, Figure S6D**), and they were significantly less enriched for IGHV1-72 compared to founders at both time points (day 25, p=0.026; day 32, p=0.016, **Figure 5E**).

**Figure 5.**
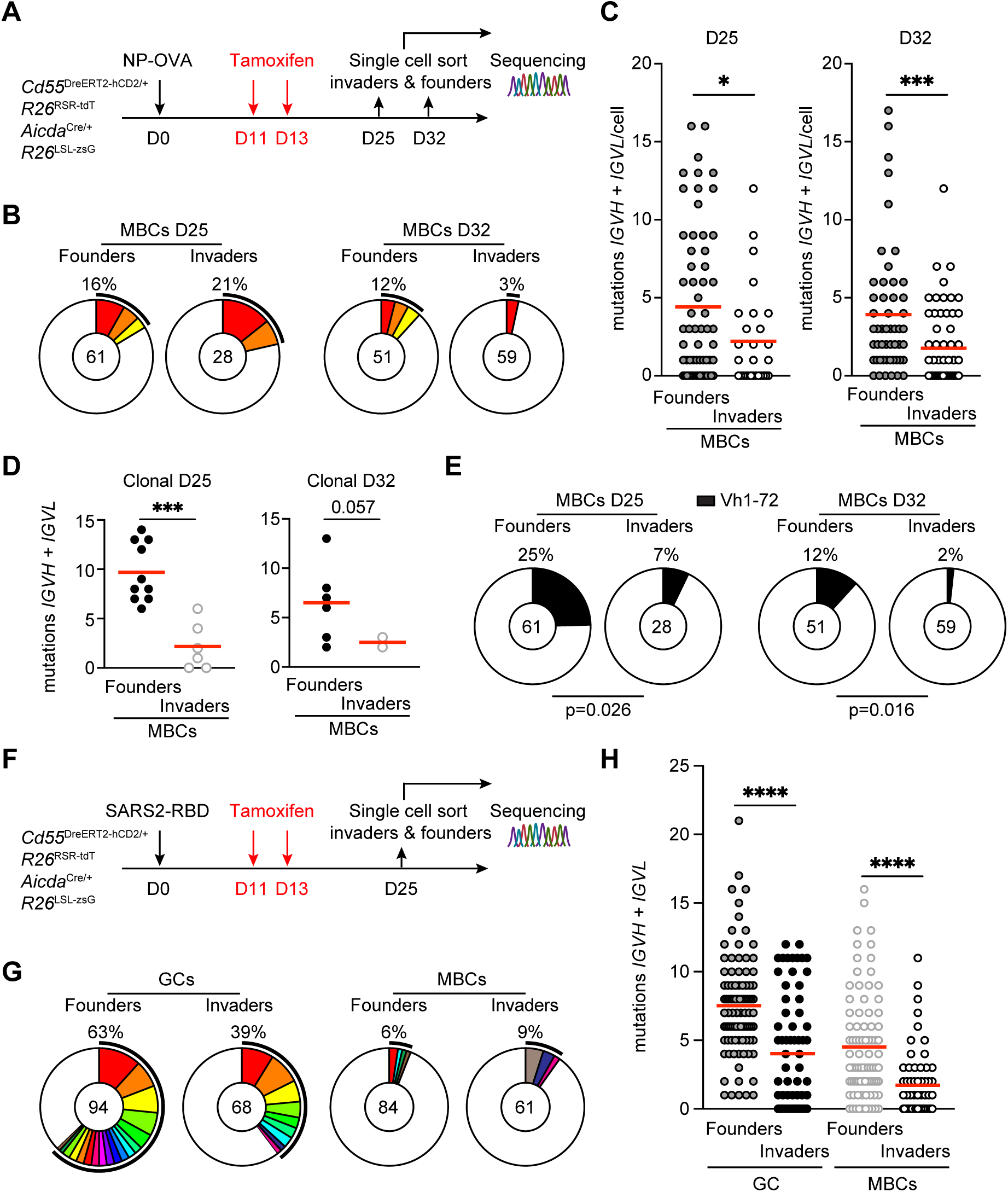
BCR repertoire and affinity of invader derived memory B cells. **(A)** Schematics of the experiment. CD55^DreER^AID^Cre^ dual reporter mice were immunized with NP-OVA, administrated with tamoxifen on day 11 and 13 and pLN from eight mice were collected on days 25 and 32 for single cell sorting and sequencing of tdT^+^ (invaders) and tdT^-^ (founders) hCD2^-^ zsG^+^ MBCs. **(B)** Pie charts depicting the distribution of antibody sequences (paired VH+VL) for Founders-MBCs (left) and Invaders-MBCs (right), pooled from 6 individual mice at days 25 and 32, respectively. The number in the inner circle indicates the number of sequences analyzed. The pie slice size is proportional to the number of clonally related sequences. The black outline and associated numbers indicate the percentage of clonal sequences detected at each time point. **(C)** Number of somatic mutations (nucleotides) of paired VH+VL chains. Each dot represents one cell with paired Ig genes. Red line indicates mean. Data were pooled from two independent experiments. Two-tailed unpaired t-test, *P ≤ 0.05, ***P ≤ 0.001. **(D)** Graph showing the number of paired VH+VL mutations of the expanded clones in Founders-MBCs and Invaders-MBCs on day 25 (left) and 32 (right). Each dot represents one cell with paired Ig genes. Two-tailed unpaired t-test, ***P ≤ 0.001. **(E)** Pie charts depicting the enrichment of IGHV1-72 clones from Ig gene sequencing analysis of Founders-MBCs and Invaders-MBCs on day 25 (left) and 32 (right). The pie slice size is proportional to the number of IGHV1-72 clones with the percentage outlined on top. The number in the inner circle indicates the number of sequences analyzed. Two-tailed Fisher’s exact test. **(F)** Schematics of the experiment. CD55^DreER^AID^Cre^ dual reporter mice were immunized with SARS2-RBD, administrated with tamoxifen on day 11 and 13 and pLN were collected on days 25 for single cell sorting and sequencing of tdT^+^ (invaders) and tdT^-^ (founders) GCs and MBCs. **(G)** Pie charts depicting the distribution of antibody sequences (paired VH+VL) for GC founders, GC invaders, Founders-MBCs and Invaders-MBCs, pooled from 8 individual mice. The number in the inner circle indicates the number of sequences analyzed. The pie slice size is proportional to the number of clonally related sequences. The black outline and associated numbers indicate the percentage of clonal sequences detected at each time point. Same colored slice between GCs (left) and MBCs (right) indicates the clones found from both compartments. **(H)** Number of somatic mutations (nucleotides) of paired VH+VL chains. Each dot represents one cell with paired Ig genes. Red line indicates mean. Data were pooled from three independent experiments. One-way ANOVA, ****P ≤ 0.0001.

To validate these findings using a vaccine antigen, we characterized tdT^+^hCD2^-^zsG^+^ GC B cells, MBCs and controls purified from dual reporter mice immunized with SARS2-RBD (**Figure 5F**). As expected, both sets of MBCs were significantly less clonal than their contemporaneous GC counterparts (**Figure 5G**). In addition, the two populations differed in that invader-derived MBCs showed fewer mutations than founder-derived MBCs (GC, p<0.0001; MBCs, p<0.0001, **Figure 5H and Figure S6E**). The data are consistent with the idea that invader origin MBCs express a less mutated group of antibodies with generally lower affinity than those derived from GC founder B cells.

### Epitope mapping

The observation that invader GC B cells successfully engage in the GC reaction and differentiate whilst expressing low affinity BCRs raises the question of how these cells remain competitive with founders in the selective environment of the GC. We hypothesized that invaders and progeny could target different antigenic sites on the RBD protein than founders. To determine whether invader derived MBC antibodies target a distinct set of antigenic epitopes, we produced 187 representative monoclonal antibodies obtained from contemporaneous founder and invader GC B cells and MBCs isolated from CD55^DreER^AID^Cre^ indicator mice immunized with SARS2-RBD (**Figure 5F**). To increase the likelihood of identifying antigen binding antibodies, we included most of the antibodies derived from expanded clones shown in **Figure 5G**. In ELISA assays (**Figure 6A**), 21/30 and 18/40 of the antibodies derived from GC B cells and 16/56 and 15/61 of those derived from founder and invader MBCs respectively showed measurable binding (**Figure 6B**).^13,27^ To map the antigenic epitopes recognized by these antibodies, we performed competition biolayer interferometry (BLI) experiments in which a preformed antibody-RBD complex comprising one of four structurally characterized antibodies^8,28,29^ was challenged with a monoclonal antibody derived from the GC or MBC compartment (**Figure 6C**). Antibodies obtained from GC founder B cells primarily targeted combinations of class-1 (57%), -2 (43%) epitopes and to a lesser extent class-3 (4%) epitopes (**Figure 6D** and **Figure S7A**). In contrast, 27% and 33% of the GC invader antibodies targeted class-1, -2 epitopes respectively, while a substantial number, 22% of these antibodies, targeted class-3 epitopes (p=0.028, **Figure 6D-E** and **Figure S7A**). In addition, 44% of invader antibodies targeted unclassified epitopes, whereas only 24% founder antibodies did so (**Figure 6D-E** and **Figure S7A)**. Similarly, founder MBCs primarily recognize class-1 -2 epitopes, while invaders are more likely to target class-3, -4 and unclassified epitopes (p=0.001, **Figure 6D-E and Figure S7A**). Taken together, there was a significant shift in the distribution of epitopes targeted by GC invaders and invader-derived MBCs compared with founder compartments (**Figure 6E**). Thus, invader GC B cells and their progeny tend to recognize subdominant epitopes that are underrepresented by the antibodies produced by their founder counterparts.

**Figure 6.**
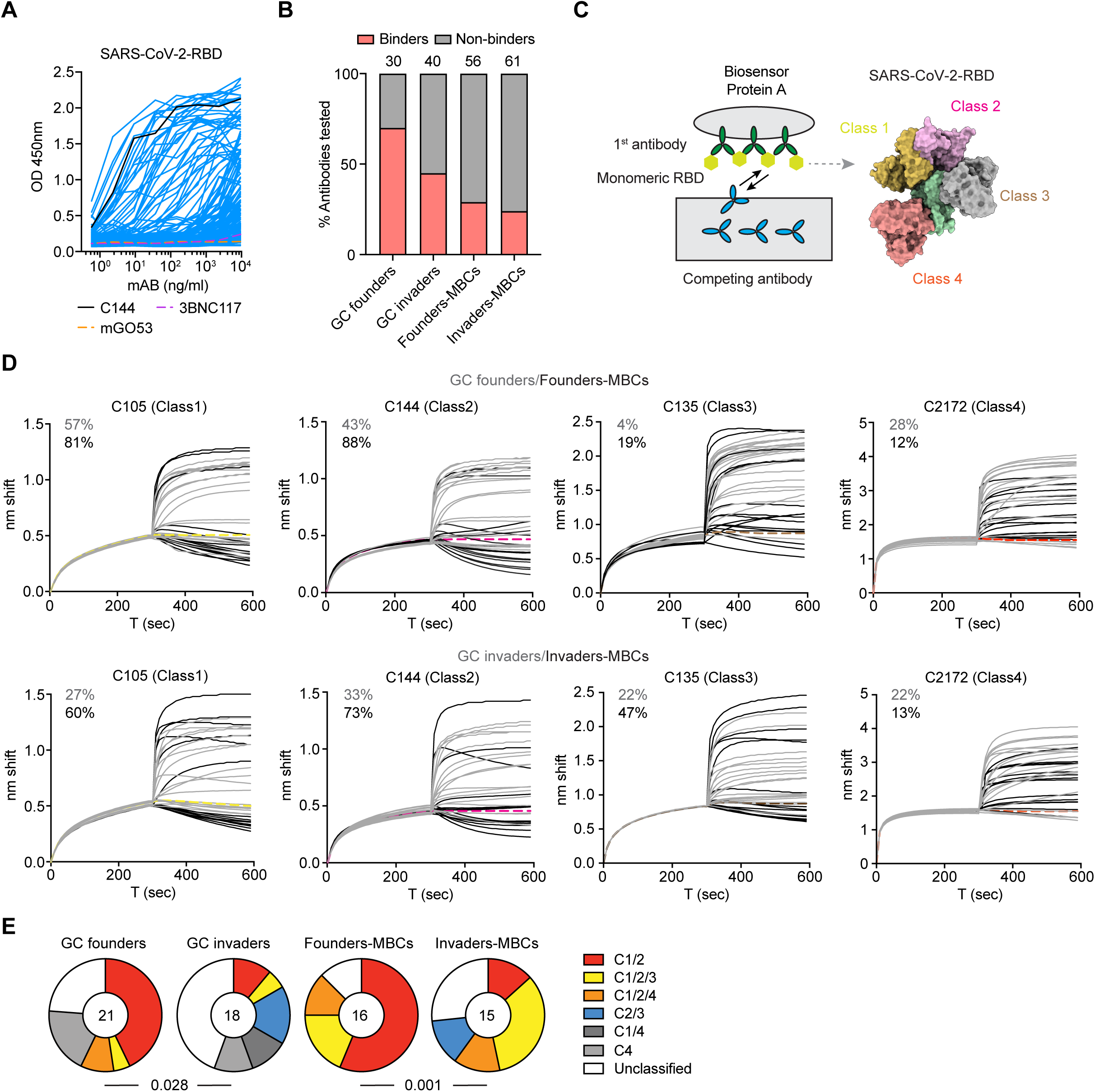
Epitope mapping. **(A)** ELISA binding of 187 monoclonal antibodies to SARS-CoV-2 RBD. Antibodies are derived from GC founders, GC invaders, Founders-MBCs and Invaders-MBCs isolated from eight CD55^DreER^AID^Cre^ dual reporter mice immunized with SARS2-RBD in Figure 5F. Each blue curve represents one antibody. C144 (black curve) antibody is the positive control, 3BNC117 and mGO53 (dash curve) are the isotype controls. **(B)** Graph showing the percentage of binding antibodies in each group. Numbers on top of each bar indicate the number of antibodies produced and tested in that group. **(C)** Schematics of the competition Biolayer interferometry (BLI) experiment. First, the capture antibody of known epitope specificity (class-reference antibody) was bound to the biosensor chip; second, the bound antibody was exposed to monovalent SARS-CoV-2 RBD antigen; and third, the antibody-RBD complex on the chip was immersed in a solution with the antibody of interest. **(D)** BLI traces obtained under monovalent competition conditions. Traces show initial association curve (antigen capture phase of the first antibody) and subsequent addition of secondary antibodies of unknown class. Thin solid grey lines represent antibodies isolated from GC founders or GC invaders. Solid black lines represent antibodies isolated from Founders-MBCs or Invaders-MBCs. Up panel shows founder antibodies. Bottom panel shows invader antibodies. Thick dashed lines are control antibodies for each class, C105 (Class1, yellow), C144 (Class2, pink), C135 (Class3, brown) and C2172 (Class4, orange) respectively. Numbers indicate the percentage of antibodies belonging to each class. **(E)**. Pie charts depicting the distribution of the epitopes targeted. The number in the center indicates the number of antibodies tested. Statistical significance was determined using a two-tailed Fisher’s exact test.

### Preferential differentiation into memory

In contrast to relatively transient GC reactions elicited by protein immunization in adjuvant,^30,31^ infections with pathogens are often associated with long-lived germinal center reactions.^1,2^ For example infection with mouse-adapted influenza A/Puerto Rico/8/1934 (PR8) elicits a potent GC reaction in the mediastinal lymph node (mLN), with GC B cells comprising 12%-20% of all B cells at days 14-42, decreasing to 4% on day 98 after-infection (**Figure S8A**). As expected, the frequency of hemagglutinin (HA) binding GC B cells decreased over time,^14,32^ suggesting GC remodeling throughout the response (**Figure S8B and C**).

To determine whether naïve B cell invasion contributes to MBC and PC production after infection, dual reporter mice were infected with PR8 and administered tamoxifen to label naïve B cells starting on days 11 and 13 after infection and bi-weekly thereafter (**Figure 7A** and **S8D-G**). To confirm that invader B cells engage with germinal center structures,^13-15,20^ we imaged the mLNs obtained on days 14 and 98 after infection. tdT^-^zsG⁺ GC cells were observed on both days 14 and 98, whereas tdT⁺zsG⁺ naïve origin cells were absent from GCs on day 14 but were abundant on day 98 (**Figure 7B and C**).

**Figure 7.**
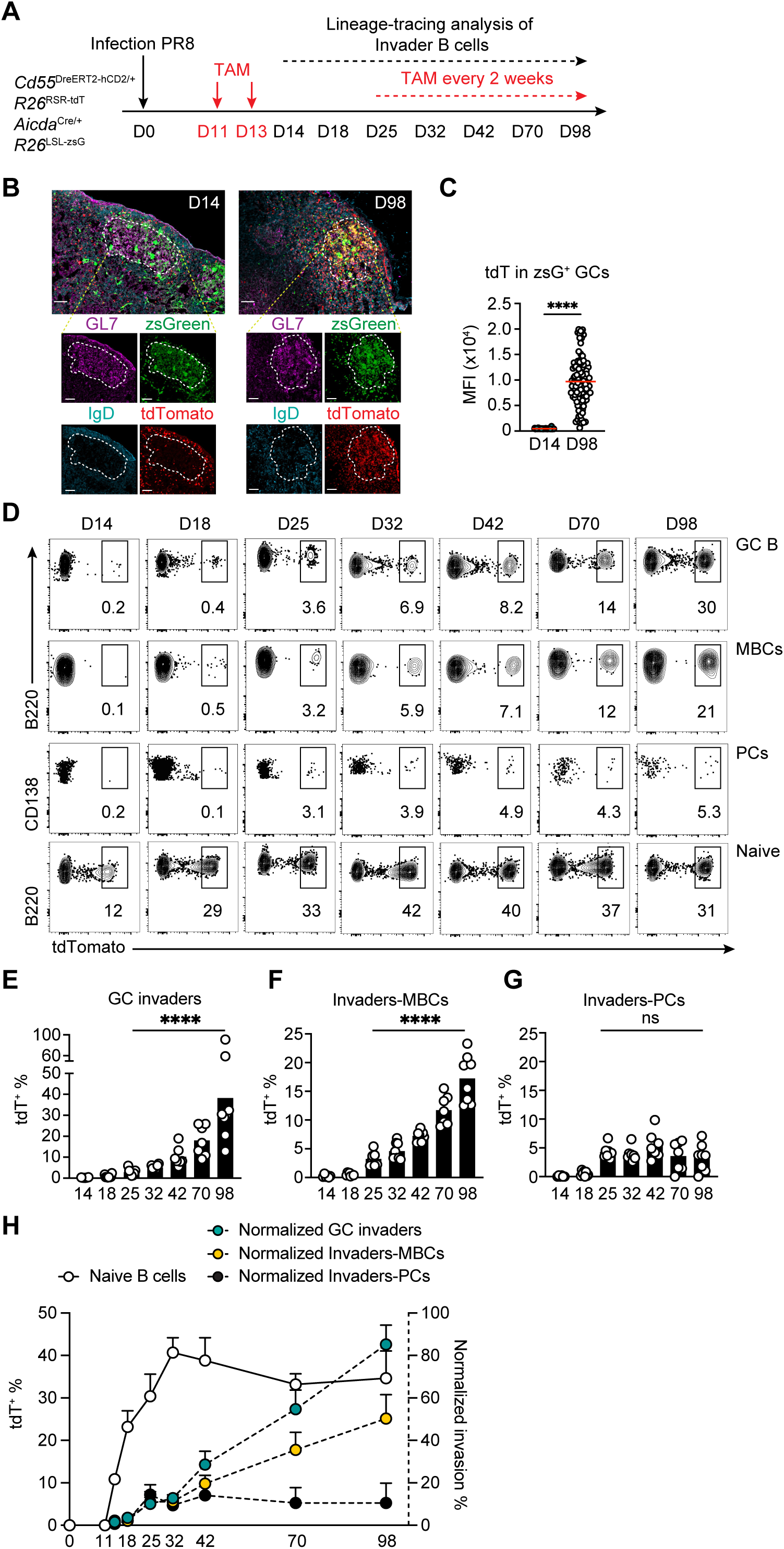
GC invaders preferentially differentiate into MBCs. **(A)** Schematics of the experiment. CD55^DreER^AID^Cre^ dual reporter mice were infected with influenza PR8 virus (30 PFU), administrated with tamoxifen on day 11 and 13 and mediastinal lymph nodes (mLN) were assayed on days 14, 18, 25, 32, 42, 70 and 98. To maintain stable labeling of naïve B cells, tamoxifen was administered orally biweekly starting on day 25. **(B)** Representative confocal microscopy images of mLNs isolated from CD55^DreER^AID^Cre^ mice 14 days or 98 days post PR8 infection. Large tiles show GC structures with inset boxes show individual channels of GL7 (purple), IgD (blue), *Aicda*^CreERT2^-*R26*^zsGreen^ (green), *Cd55*^DreERT2-hCD2/+^-*R26*^tdTomato^ (red). Scale bar = 50um. **(C)** Statistics of the mean fluorescence intensity (MFI) of tdTomato in zsGreen⁺ germinal center (GC) B cells. Each dot represents a single zsGreen^+^ GC B cell, with data collected from 4 mice at each time point (day 14 and day 98). Red line indicates mean, two-tailed unpaired t-test, ****P ≤ 0.0001. **(D)** Representative flow cytometry plots summarizing the percentage of tdT^+^ GCs, tdT^+^hCD2^-^zsG^+^ MBCs, tdT^+^zsG^+^ PCs and tdT^+^ naïve B cells on days 14, 18, 25, 32, 42, 70 and 98. **(E-G)** Statistics of tdT^+^ GC B cells **(E)**, tdT^+^ MBCs **(F)** and tdT^+^ PCs **(G)** show in **(D)**. Each dot represents one mouse, bar indicates mean, two-tailed unpaired t-tests between day 25 and day 98, ****P ≤ 0.0001. **(H)**. Graph showing the percentage of tdT^+^ naïve B cells (white) over time. Green circles show frequency of tdT^+^ GC B cells, yellow circles show frequency of tdT^+^hCD2^-^zsG^+^ MBCs and black circles show tdT^+^zsG^+^ PCs normalized for the labeling efficiency of the naïve B cell compartment. 3 independent experiments, circle indicates mean, and error bars indicate SD, n = 6-8 at each time point.

The contribution of late entering naive B cells to GC B cell, MBC and PC compartments was assessed by flow cytometry on days 14, 18, 25, 32, 42, 70, and 98 after infection. Naïve B cell labeling peaked at approximately 41% at day 32 remaining relatively stable thereafter (**Figure 7D and Figure S8H)**. In contrast, tdT^+^hCD2^-^zsG^+^ invader cells were barely detectable in the GC until day 25 but increased to 30% of all GC B cells on day 98 or an average of 85% after correcting for labelling efficiency (**Figures 7D-H)**. MBC derived from naïve B cells entering late increased to 50% of all MBCs after correcting for labeling by naïve B cells on day 98 (**Figures 7D-H, Figure S8I**). In contrast, differentiation of invaders into PCs was limited to 9-14% after correction for labeling efficiency between days 25-98 (**Figure 7G and H**). To examine the contributions of MBC and PC production versus egress we performed pulse-chase experiments (**Figure S9A-C**). The frequency and number of tdT^+^hCD2^-^zsG^+^ MBCs increased progressively over time, whereas the frequency and number of tdT^+^zsG^+^ PCs remained relatively stable (**Figure S9D-E**). Notably, FTY720 treatment did not affect the frequencies of tdT^+^ cells in either population, indicating that the preferential differentiation of invader cells into MBCs rather than PCs reflects a difference in production rather than egress. Thus, GC invader B cells preferentially give rise to increasing numbers of MBCs over time.

## Discussion

Naïve B cells are continually recruited to enter GCs throughout the immune response.^13-15^ GC invaders express low affinity receptors for the initiating immunogen and are typically less somatically mutated, reflecting less time spent in the GC, than their founder counterparts. Here we report on the development of dual reporter CD55^DreER^AID^Cre^ indicator mice that enabled fate-mapping naïve B cells entering the GC in the late stages of the response and their development into PCs and MBCs. GC invaders preferentially give rise to MBCs that harbor fewer somatic mutations, display lower affinity, and target subdominant epitopes relative to founder clones. Thus, the antibodies they produce diversify the memory compartment.

PCs and MBCs originating in GCs shape long-term humoral immunity. Our lineage-tracing experiments indicate that PC production parallels GC size peaking after approximately 2 weeks and decreasing thereafter while MBC production increases in proportion to the GC. This observation is consistent with recently documented similarities between GC B cell and PC selection.^33^ GC light zone cells and PC precursors are selected to divide in direct proportion to the strength of T_FH_ signaling and the amount of c-Myc they produce.^33-37^ At the peak of the GC, abundant antigen enables higher levels of antigen capture and peptide-MHC presentation to T_FH_ by GC light zone cells and PC precursors and subsequently greater levels of cell division and clonal expansion.^1,2^ In contrast, relatively lower levels of antigen capture and presentation favor MBC development leading to a proportional increase in cells being selected into this compartment over time.^16,27^ The overall kinetics of PC and MBC production from our experiments suggest the requirement for stronger T_FH_ signaling to produce PCs and weaker for MBCs.

As originally noted by Landsteiner and Eisen, the frequency of antibodies targeting subdominant epitopes increases with time.^3,4^ Several mechanisms appear to contribute to this shift including epitope masking by antibodies,^38,39^ and increasing levels of T cell help.^40,41^ Epitope masking prevents recruitment and subsequent selection of cells that target initially dominant epitopes that rapidly elicit high titers of antibodies.^8,42^ For example in SARS-CoV-2 infection or vaccination, the antibodies that are initially produced target highly exposed Class 1 and 2 epitopes.^8,42^ These antibodies block their epitopes favoring antibody evolution to more conserved subdominant epitopes over time.^38^ Increasing antigen valency in immune complexes lowers affinity thresholds for B cell selection by avidity effects and increasing levels of T cell help decrease the competitive advantage of high affinity cells that present more peptide-MHC. These phenomena are consistent with our observation that the affinity threshold for GC entry decreases over time^13^ and that invaders produce lower affinity MBCs targeting subdominant epitopes.

Antibody diversity is a key feature of polyclonal humoral immune responses. Beginning in the bone marrow with V(D)J recombination, and subsequent somatic mutation in GCs, genome-based DNA transactions ensure the existence of antibodies that can target a very broad range of potential pathogens. Production of an affinity diverse cohort of MBCs that target both dominant and subdominant antigenic epitopes in GCs enhances diversity and safeguards the immune systems’ ability to defend against evolving pathogens. Subdominant epitopes such as the influenza hemagglutinin stalk or the HIV Env CD4-binding site may be more conserved between strains or better targets for neutralization than immunodominant epitopes. Our data indicate that naïve B cells entering GCs at later stages of the reaction contribute to the diversification of the MBC compartment.

In conclusion, GC output is dynamically modulated over the course of the immune response in a way that coincides with potential pathogen evolution. Late-arriving invader B cells contribute to diversity by differentiating into MBCs targeting suboptimal epitopes that likely bolster protection against rapidly evolving pathogens including SARS-CoV-2, and influenza. However, immune evolution away from a singular epitope may interfere with the development of immune responses that require specific epitope targeting as in HIV-1 sequential vaccine approaches.

## Limitation of the study

Invader cells preferentially give rise to MBCs that recognize subdominant epitopes, and it would be intriguing to investigate how these cells behave in comparison to naïve and founder-derived MBCs during a secondary response. However, the current CD55^DreER^AID^Cre^ dual-reporter system does not permit recall experiments in the same mouse. Because AID expression is constitutive, the dual reporter cannot distinguish between GC B cells participating in the primary versus secondary responses. The same limitation applies when assessing the extent to which MBCs may re-enter an ongoing primary GC reaction. In addition, whereas the *S1pr2*^CreERT2^ and AID^Cre^ reporters efficiently label GC B cells, they may also mark a fraction of extrafollicularly activated B cells throughout the response.^22^ The future development of genetic or alternative strategies that can exclusively track GC-derived progeny and enable the evaluation of invader-derived MBCs during recall responses would be of great interest.

## Supplementary Figure legends

**Figure S1. Temporal dynamics of germinal center output, related Figure 1.**

**(A)** Gating strategy for zsGreen^+^ GC B cells, MBCs and PCs. Representative plots are from popliteal lymph nodes of *S1pr2*^CreERT2/+^ *Rosa26*^LSL-zsGreen/+^ mice immunized with NP-OVA and treated with tamoxifen. **(B)** Statistics showing the number of circulating B220⁺ B cells in the blood before and three days after FTY720 treatment in *S1pr2*^CreERT2/+^ *Rosa26*^LSL-zsGreen/+^ mice immunized with NP-OVA and treated with tamoxifen. Bar indicates mean. P values calculated using Mann-Whitney, ****P ≤ 0.0001.

**Figure S2. Generation of GC invader fate-mapping mice, related Figure 2.**

**(A)** Conditional labeling of follicular B cells in *Cd55*^CreERT2/+^ *Rosa26*^LSL-tdTomato/+^ mice. Mesenteric lymph nodes were analyzed 4 days after tamoxifen administration in naïve mice. Labeled tdT^+^ cells in B220^+^ CD38^+^ Fas^-^ non-GC B cells, B220^+^ CD38^-^ Fas^+^ GC B cells and B220^lo^ CD138^+^ plasma cells were analyzed by flow cytometry. Left, representative flow cytometry plot. Right, statistics of tdTomato^+^ cells. **(B)** Examination of CD55 expression in *Aicda*^Cre/+^ *Rosa26*^LSL-zsGreen/+^ mice. MBCs (B220^+^ CD38^+^ Fas^-^ zsGreen^+^) and naïve B cells (B220^+^ CD38^+^ Fas^-^ zsGreen^-^) in mesenteric lymph nodes were analyzed. Left, representative flow cytometry plot. Right, statistics of statistics of CD55 GMFI. **(C)** Targeting strategy and the configuration of the *Cd55*^L-DreERT2-hCD2-L/+^ allele. The mouse strain was produced in Nussenzweig Lab at The Rockefeller University and crossed to *Rosa26*^RSR-tdTomato^ to create *Cd55*^DreERT2-hCD2^ *Rosa26*^RSR-tdTomato^ conditional indicator mice. **(D)** Representative flow cytometry plot showing the hCD2 co-expression with CD55 in knock-in and wild-type control mice. **(E)** Statistics of hCD2 expression. **(F)** Representative flow cytometry plot showing GCs and PCs in popliteal lymph nodes obtained from CD55^DreER^AID^Cre^ mice 14 days after or without immunization. **(G)** Graph showing the percentage of GC B cells in **(F). (H)** Graph showing the cell numbers of PCs in **(F).** Each dot represents one mouse, bar indicates mean, two-tailed unpaired t-test, **P ≤ 0.01.

**Figure S3. PCs development from GC invaders, related to Figure 2.**

**(A-B)** Representative flow cytometry plot showing GCs and PCs in popliteal lymph nodes obtained from CD55^DreER^AID^Cre^ dual reporter mice 14, 18, 25 or 32 days after immunization with NP-OVA. **(C)** Statistics of GCs and PCs shown in **(A-B)** between day 14 and day 32. **(D)** Representative flow cytometry plot showing the hCD2 expression among zsGreen labeled PCs and the gating of the tdTomato labeled PCs assessed in **(B)**. **(E)** Schematics of the experiment. CD55^DreER^AID^Cre^ dual reporter mice were immunized with Tenivac, administrated with tamoxifen on day 11 and 13 and popliteal lymph nodes were assayed on days 14, 18, 25, and 32. **(F)** Graphs showing the percentage of tdTomato labeled GCs and PCs at each time points. Each dot represents one mouse, bar indicates mean, two experiments. One-way ANOVA, *P ≤ 0.05, **P ≤ 0.01, ***P ≤ 0.001, ****P ≤ 0.0001.

**Figure S4. Confocal imaging of pLN from dual reporter mice, related Figure 3.**

**(A)** Schematic representation of the experimental protocol. Three CD55^DreER^AID^Cre^ dual reporter mice were immunized with NP-OVA, administrated tamoxifen on day 11 and popliteal lymph nodes (pLN) imaged on day 15. Three additional dual reporter mice were immunized with NP-OVA, administrated tamoxifen on day 0 and popliteal lymph nodes (pLN) imaged on day 4. **(B-C)** Representative confocal microscopy images of pLNs isolated from CD55^DreER^AID^Cre^ mice 15 days or 4 days post NP-OVA immunization. Large tiles show whole lymph node structures with inset boxes show individual channels of GL7 (purple), IgD (blue), *Aicda*^CreERT2^-*R26*^zsGreen^ (green), *Cd55*^DreERT2-hCD2/+^-*R26*^tdTomato^ (red). Scale bar = 100um. **(D)** Left, quantification of zsG^+^tdT^+^ cells in the pLN at day 15 or day 4. Right, graph showing the percentage of zsG^+^tdT^+^ cells within the follicular area (Fo) or non-follicular (Non-Fo) region of pLN at day 15 or day 4.

**Figure S5. BCR sequencing of invader-derived PCs, related to Figure 3.**

**(A)** Equation to calculate purity of founder control cells. **(B)** Graph showing the calculated purity of founder cells in GCs obtained from CD55^DreER^AID^Cre^ dual reporter mice 25 days after immunization and 14 days after tamoxifen administration. **(C)** Pie charts depicting the distribution of antibody sequences (paired VH+VL) for GC founders, GC invaders, Founders-PCs and Invaders-PCs from 6 individual mice. The number in the inner circle indicates the number of sequences analyzed. The pie slice size is proportional to the number of clonally related sequences. The black outline and associated numbers indicate the percentage of clonal sequences detected at each time point. **(D)** Graph shows EC_50_ of Fabs binding to NP_26_-BSA. Each dot represents one Fab. Red line indicates median. One-way ANOVA, *P ≤ 0.05. **(E)** IGHV1-72 usage among NP-binding Fabs within each compartment. Two-tailed Fisher’s exact test.

**Figure S6. Memory B cells, related to Figure 4 and Figure 5.**

**(A)** Representative flow cytometry plot showing MBCs in popliteal lymph nodes obtained from CD55^DreER^AID^Cre^ dual reporter mice 14 days after or without immunization. **(B)** Graph showing the cell numbers of MBCs in **(A)**. Each dot represents one mouse, bar indicates mean, two experiments. two-tailed unpaired t-test, ****P ≤ 0.0001. **(C)** Numbers of tdTomato^+^ hCD2^-^zsG^+^ MBCs (White circles) and tdTomato^+^ zsG^+^ PCs (Black, Invaders-PCs) in CD55^DreER^AID^Cre^ dual reporter mice after NP-OVA immunization. **(D)** Graph showing the distribution frequency of somatic mutations in Founders-MBCs (left, green) and Invaders-MBCs (right, yellow) isolated from day 25 (up) and 32 (bottom) respectively from CD55^DreER^AID^Cre^ dual reporter mice immunized with NP-OVA and injected with tamoxifen on day 11 and 13. **(E)** Graph showing the distribution frequency of somatic mutations in Founders-MBCs (green) and Invaders-MBCs (yellow) isolated on day 25 from CD55^DreER^AID^Cre^ mice immunized with SARS2-RBD and injected with tamoxifen on day 11 and 13.

**Figure S7. Antibody epitope mapping, related to Figure 6.**

Heatmap showing the relative inhibition of secondary testing antibody binding to preformed capture antibody-RBD complexes (grey indicates no binding, red indicates binding, and pink indicates indeterminate results).

**Figure S8. Longitudinal analysis of GC invaders contributing to MBCs and PCs, related to Figure 7.**

**(A)** Graph showing the percentage of GCs over time in mediastinal lymph nodes (mLN) obtained from CD55^DreER^AID^Cre^ dual reporter mice infected with PR8. **(B)** Representative flow cytometry plot showing influenza PR8 hemagglutinin (HA)-bait binding cells in GCs and in naïve B cell controls in mLN obtained from CD55^DreER^AID^Cre^ dual reporter mice infected with PR8. **(C)** Graph showing the percentage of HA-bait binding GC B cells over time. Each dot represents one mouse, bar indicates mean. **(D)** Representative flow cytometry plot showing the gating strategy for the longitudinal tracking of invaders in GCs, MBCs and PCs. **(E-G)** Graph showing the percentage of GC B cells, percentage of hCD2⁻ MBCs, and PCs over time. **(H)** Graph showing the tdTomato labeled naïve B cells over time. Each dot represents one mouse, bar indicates mean. **(I)** Graph showing the frequency of tdTomato-labeled naïve B cells and hCD2⁻ MBCs in mediastinal lymph nodes from the uninfected CD55^DreER^AID^Cre^ dual reporter mice. Tamoxifen was provided biweekly. Red circles show mean frequency of tdT^+^ naïve B cells and white circles indicate the mean frequency of tdT^+^zsG^+^hCD2⁻ MBCs.

**Figure S9. Pulse-chase experiments to assess the generation of invader-progeny and their egress from lymph nodes, related Figure 7.**

**(A)** Schematic representation of the experimental protocol. CD55^DreER^AID^Cre^ dual reporter mice were infected with influenza PR8 virus (30 PFU), and a single tamoxifen pulse was administered on days 18, or 25, or 35 post infection to label naïve B cells. Cell egress from draining lymph node was prevent by two FTY720 injections as indicated, and GC B cells, MBCs and plasma cells were analyzed on days 25, or 32 or 42. **(B)** A parallel pulse-chase experiment was conducted without FTY720 treatment. **(C)** Representative flow cytometry plots showing the gating of tdT^+^ cells among MBCs and PCs. MBCs were gated on hCD2^-^zsG^+^, and PCs were zsG^+^. **(D-E)** Bar graphs showing the normalized percentages after correcting for labeling efficiency of the naïve B cell compartment **(D)**, or numbers **(E)** of tdT^+^ cells in MBCs (black bar) or PCs (white bar) at each time point. Each dot represents one mouse.

## ACKNOWLEDGMENTS

We thank T. Eisenreich for help with mouse colony management, Masa Jankovic and T. Waldetario for laboratory support, K. Gordon and J-P. Truman for cell sorting, all Nussenzweig laboratory members for discussion. We thank T. Kurosaki for S1PR2 mice. This work was supported in part by NIH Center for HIV/AIDS Vaccine Immunology and Immunogen Discovery (CHAVID) 1UM1AI144462-01 to M.C.N., and the Stavros Niarchos Foundation Institute for Global Infectious Disease Research. This article is subject to HHMI’s Open Access to Publications policy. HHMI lab heads have previously granted a non-exclusive CC BY 4.0 license to the public and a sublicensable license to HHMI in their research articles. Pursuant to those licenses, the author-accepted manuscript of this article can be made freely available under a CC BY 4.0 license immediately upon publication. M.C.N. is a Howard Hughes Medical Institute (HHMI) investigator.

## AUTHOR CONTRIBUTIONS

P.Z. and M.C.N. conceived the study, designed experiments, and interpreted data. P.Z. designed and generated the CD55^DreER^ mice with the help from H.H., and P.Z., H.H., A.J.M., and V.R. performed experiments. A.G., K.Y., Z.W. and B.H. provided critical reagents needed to perform experiments. P.Z. and M.C.N. wrote the manuscript with input from all co-authors.

## EXPERIMENTAL MODEL AND SUBJECT DETAILS

### Mice

C57BL/6J (stock No 000664), *Rosa26*^LSL-zsGreen^ (Ai6, Rosa-CAG-LSL-ZsGreen1-WPRE^43^, stock No 007906), and *Rosa26*^LSL-tdTomato^ (Ai14, Rosa-CAG-LSL-tdTomato-WPRE^43^, stock No 007914) mice were obtained from Jackson Laboratories. *S1pr2*^CreERT2^ transgenic mice were generated and kindly provided by T. Kurosaki.^16^ *Rosa26*^RSR-tdTomato^ (Ai66R, Rosa-CAG-Rox-STOP-Rox-tdTomato^44^) mice were sourced from D. Mucida Lab. *Cd55*^CreERT2^ mice^13^ and *Aicda*^Cre^ mice^21^ were generated and maintained at the Rockefeller University. *Cd55*^loxp-IRES-DreERT2-hCD2-loxp^ mice were generated by CRISPR/Cas9 editing and recombinant Adeno-associated viruses serotype 6 (rAAV6) infection of C57BL/6J mouse embryos with details described in the methods. Female and male mice were used in this study and were 6 weeks of age or older at the time of any experimentation. All mice were bred, maintained, and immunized under SPF (specific pathogen-free) conditions at 22 °C and 30–70% humidity, following a 12-hour light/dark cycle with ad libitum access to food and water, at the Rockefeller University Comparative Biosciences Center. All animal procedures were performed in accordance with protocols approved by the Rockefeller University Institutional Animal Care and Use Committee (IACUC).

## METHOD DETAILS

### Generation of *Cd55*^loxp-IRES-DreERT2-hCD2-loxp^ mice by CRISPR/Cas9 and AAV viruses

We designed the allele as indicated in Figure S2. Template DNA fragments were purchased from IDT and cloned into the pAAV6 vector in between AAV2 inverted terminal repeats by Gibson assembly. 293AAV cell line (CellBiolabs, AAV-100) were co-transfected with pAAV vector carrying template insertion and TESSA-RepCap6 vector as previously described.^45^ Transfected cells were incubated for 4 days and recombinant AAV6 virus were collected in PEG/NaCl solution. rAAV6 was extracted by Chloroform (Sigma-Aldrich 650471-1L) after PEG/NaCl precipitation and buffer-exchanged with Formulation Buffer (DPBS with 0.001% Pluronic F68, Gibco 24040-032). rAAV6 virus titer was quantified using qPCR. 15-25 mouse embryos were incubated with around 2 x 10^9^ genome copies (GC) of rAAV6 for 6 hours at 37 °C. Cas9/sgRNA complexes targeting the *Cd55* locus were electroporated into mouse embryos, which were then recovered and incubated overnight at 37 °C before implantation into female mice. We verified correct allele insertion by Sanger sequencing across the entire locus using genomic primers located outside the homology arms. The founder mouse was backcrossed for at least five generations onto C57BL/6J mice before being used in experiments.

### Immunizations and treatments

Immunizations were performed at the footpad with 50 uL of PBS containing 25 ug of NP-OVA (Biosearch technologies, NP conjugation ratio 17) precipitated in aluminum hydroxide gel adjuvant (Alhydrogel, Invivogen, vac-alu-250) at a 2:1 ratio. 2, 3 and 4 μg of SARS-CoV-2 RBD were used to immunize the mice in an escalating dose format on day 1, 3 and 5, with Alhydrogel adjuvant only accompanying the protein on the first injection. 50ul of Tenivac^®^ (tetanus and diphtheria toxoids, adsorbed; Sanofi Pasteur) was administered per footpad.

Activation of the Cre or Dre recombinase in the *S1pr2*^CreERT2^, *Cd55*^CreERT2^, and *Cd55*^DreERT2-hCD2^ mice was induced by oral administration of 12 mg tamoxifen (Sigma, T5648) in 200 µL of corn oil (Sigma, C8267) at the indicated time points.

To prevent cell egress from LNs, 1mg/kg FTY720 (Selleck Chemicals) was administered intravenously at the indicated time points.

### Influenza infection and fate-mapping

Mice were anesthetized with isoflurane (2%), after which 30 PFU of mouse-adapted influenza PR8 virus (provided by J. Ravetch lab, Rockefeller University) were intranasally injected. Fate-mapping of GC invaders in CD55^DreER^AID^Cre^ mice was carried out by oral administration of 12 mg tamoxifen (Sigma, T5648) in 200 µL of corn oil (Sigma, C8267) at the indicated time points to activate and maintain the expression of Dre recombinase. At different time points after the infection, the draining mediastinal lymph nodes were harvested for various readouts. All infected mice were held at the designated BSL2 room at the Rockefeller University Comparative for Biosciences Center.

### Protein expression and bait preparation

The mammalian expression vector encoding the RBD of SARS-CoV-2 (GenBank MN985325.1; S protein residues 319–539) for immunization and BLI experiments was previously described.^29^ Vectors used for recombinant HA trimer protein were kindly provided by J. Ravetch lab (Rockefeller University) and P. Wilson lab (Weill Cornell Medical College). All constructs were confirmed by Sanger sequencing and used to express soluble proteins via transient transfection of Expi293F cells (Gibco/Thermo Fisher Scientific, A14527). Supernatants were collected after 4 days, and the RBD or HA proteins were purified by nickel affinity chromatography. Peak fractions from size-exclusion chromatography were identified using native gel electrophoresis, and the fractions corresponding to monomeric RBDs or the HA trimer were pooled and stored at -20 °C.

Purified HA trimer protein was biotinylated using the EZ-Link-Sulfo-NHS-LC-Biotinylation kit according to the manufacturer’s instructions (Thermo Fisher Scientific, 31497). Excess biotin was removed by diafiltration with 100 kDa cutoff. Biotinylated HA-trimer was conjugated to streptavidin-AF647 (BioLegend, 405237) and to streptavidin-BV421 (BioLegend, 405225).

### Flow cytometry

LNs were collected into 1.5 ml Eppendorf tubes with FACS buffer (PBS, 2% FCS) and dissociated using a pestle. After incubation with 5 mg/mL of anti-CD16/32 (rat mAb 2.4G2, mouse Fc block BD) for 15 min at 4°C, cell surface antigens were stained for 30 min at 4°C.

BD FACSymphony was used for flow cytometric analysis. Antibodies were used as follows: Anti-mouse FC block (2.4G2, BD, 553142), Anti-CD95-PEcy7 (Jo2, BD, 557653), Anti-CD95-BUV737 (Jo2, BD, 741763), Anti-B220-BV786 (RA3-6B2, BD, 563894), Anti-TACI-BV421 (8F10, BD, 742840), Anti-CD138-BV605 (218-2, Biolegend, 142516), Anti-T-and B-cell activation antigen-PeCy7 (GL7, Biolegend, 144620), Anti-CD55-APC (RIKO-3, Biolegend, 131812), Anti-CD38-AF700 (90, ThermoFisher, 56-0381-82), Anti-CD4-eF780 (RM4-5, Invitrogen, 47-0042-82), Anti-CD8-eF780 (53-6.7, Invitrogen, 47-0081-82), Anti-NK1.1-eF780 (PK136, Invitrogen, 47-5941-82), Anti-F4/80-eF780 (BM8, Invitrogen, 47-4801-82), Anti-human CD2-PE-Vio® 770 (REA972, Miltenyi Biotec, 130-116-151), Anti-human CD2-APC (REA972, Miltenyi Biotec, 130-116-150). Live/dead marker Zombie NIR (Biolegend, 423106, and from eBiosciences).

### Confocal microscopy and imaging

Lymph nodes were prepared for confocal imaging as follows. Briefly, LNs were collected from mice and fixed in 4% paraformaldehyde (PFA) for 2 hours. The LNs were then washed in PBS, immersed in 30% sucrose for 12 hours for cryoprotection, and subsequently embedded in OCT (Optimal Cutting Temperature compound; TissueTek) blocks. Tissues were sectioned at 10 μm on a cryostat (Microm) and blocked with Fc block for 30 minutes. Slides were incubated at room temperature with the indicated antibodies GL7 (GL7, Biolegend, 144606) and IgD (11-26c.2a, Biolegend, 405725) for 6-12 hours, washed in PBS, and mounted using Fluoromount-G (Thermo Scientific). Microscopy was performed on an inverted laser scanning microscope (LSM980, Zeiss). For quantification, the Spots function in Imaris imaging software (Bitplane) was used to identify zsGreen^+^ germinal center B cells (GCs) based on zsGreen fluorescence intensity and cell quality (a metric automatically quantified by the software). These cells were further examined for mean fluorescence intensity (MFI) of tdTomato, and all spots were manually verified.

### qPCR

rAAV6 genomic copies were quantified using qPCR as previously described.^45^ In brief, rAAV6 virus were digested with DNAse I for 30 min at 37°C. Serial dilutions were measured by qPCR using forward ITR primer, 5’-GGAACCCCTAGTGATGGAGTT; and reverse ITR primer, 5’-CGGCCTCAGTGAGCGA. Full capsid AAV6 reference standard from Vigene RS-AAV6-FL was used to create standard curve. PowerUp™ SYBR™ Green Master Mix (Thermo A25742) was used to prepare the reaction. All qPCRs were performed in a 384-well plate format using the Applied Biosystem QuantStudio 6 or 7 Flex real-time PCR system.

### Single-cell index sorting and RNA purification

Single mouse GC B cells, PCs and MBCs from lymph node were stained, single cell sorted with a BD FACSymphony S6 sorter into 96-well plates containing 5 mL lysis buffer (TCL buffer (Qiagen, 1031576), and 1% 2-mercaptoethanol (Sigma, M3148) and immediately frozen at -80°C. Single-cell RNA was purified using magnetic beads (RNAClean XP, Beckman Coulter, A63987) according to the manufacturer’s instructions. The RNA was eluted from the beads with 11 uL of a solution containing random primers (14.5 ng/uL, Invitrogen, 48190-011), tergitol (0.5% NP-40 70% in H₂O, Sigma-Aldrich, NP40S), and RNase inhibitor (0.6 U/uL, Promega, N2615) in nuclease-free water (Qiagen), and then incubated at 65 °C for 3 minutes. Subsequently, cDNA was synthesized by reverse transcription using 7 uL of a solution containing SuperScript III Reverse Transcriptase, 5× buffer, dNTPs (25 mM), DTT (SuperScript III Reverse Transcriptase, Invitrogen, 18080-044, 10,000 U), and RNase inhibitor (0.6 U/uL, Promega, N2615) in nuclease-free water (Qiagen). The reaction was incubated sequentially at 42 °C for 10 minutes, 25 °C for 10 minutes, 50 °C for 60 minutes, and 94 °C for 5 minutes. The resulting cDNA was either stored at –20 °C or immediately used for antibody gene amplification by nested polymerase chain reaction (PCR) after the addition of 10 uL of nuclease-free water.

### Antibody sequencing and cloning

#### Mouse antibody genes were amplified by nested PCR

42ul of a reaction mixture containing HotStarTaq DNA polymerase (250 U/50 μL), 10× buffer (Qiagen, 203209), dNTPs (25 uM), 5′ forward primers (50 uM), and 3′ reverse primers (50 uM), along with 4 uL of cDNA (for PCR1) or PCR1 product (for PCR2) in nuclease-free water (Qiagen), were used for amplification.^46^ For the first PCR, both heavy chains (IgM and IgG) and light chains (IgK and IgL) were amplified using the following protocol: 95 °C for 15 minutes, followed by 50 cycles of 94 °C for 30 seconds, 46 °C for 30 seconds, and 72 °C for 55 seconds, with a final extension at 72 °C for 10 minutes. For the second PCR, the κ light chain was amplified using the same thermal cycling conditions as the first PCR, while the heavy chain was amplified with an adjusted annealing temperature of 55 °C, and λ light chain of 57 °C.

### Single cell VDJ Sequencing and analysis

The PCR products of antibody heavy chain and light chain genes were purified and Sanger-sequenced (Genewiz). Sequencing primers are listed as follow. Heave chain with mixed primers, AGGGGGAAGACATTTGGGAAGGAC & GAYATTGTGMTSACMCARWCTMCA. Kappa light chain with primer TGGGAAGATGGATACAGTT. Lambda light chain with primer CTCYTCAGRGGAAGGTGGRAACA. Sequencing files were analyzed using our Ig analysis pipeline. ^47^ (https://github.com/stratust/igpipeline/tree/igpipeline2_timepoint_v2).

### Antibody and Fab cloning

V(D)J sequences were synthesized as eBlocks (IDT), which included short homologies at both ends for Gibson assembly, and subsequently cloned into human IgG1 Antibody or Fab, human IgK, or human IgL2 expression vectors using the NEB Hifi DNA Assembly mix (NEB, E2621L). Plasmid sequences were confirmed by Sanger sequencing (Genewiz).

### Antibody and Fab expression

His6-tagged antibodies or Fabs, along with κ and λ light chains, were expressed by transient transfection of both heavy chain and light chain plasmids in Expi293F cells (Thermo Fisher Scientific). After buffer exchange in PBS, the Abs and Fabs were purified using Ni Sepharose 6 Fast flowresin (Cytiva) and yield productivity was checked by measurement with Nanodrop and PAGE analysis.

## ELISA

ELISA assays to measure Fab binding to NP-BSA or antibody binding to SARS-CoV-2 Wuhan-Hu-1 RBD were performed by coating high-binding 96-half-well plates (Corning, 3690) with 50 μL per well of a 1 μg/mL protein solution in PBS overnight at 4 °C. The plates were then washed six times with washing buffer (1× PBS with 0.05% Tween-20, Sigma-Aldrich) and incubated with 170 μL of blocking buffer per well (1× PBS with 2% BSA and 0.05% Tween-20, Sigma-Aldrich) for 1 hour at room temperature. Immediately after blocking, the test Fabs or antibodies were diluted in PBS and incubated for 1 hour at room temperature, starting at 10 μg/mL with 8-11 additional four or five-fold serial dilutions. After washing the plates six times with washing buffer, they were incubated with an anti-human Fab IgG secondary antibody conjugated to horseradish peroxidase (HRP) (Jackson Immuno Research, 109-036-088) in blocking buffer at a 1:5,000 dilution. The plates were developed by adding the HRP substrate 3,3′,5,5′-tetramethylbenzidine (Thermo Fisher Scientific) for 3 minutes, and the reaction was stopped by adding 50 μL of 1 M H₂SO₄. Absorbance was immediately measured at 450 nm using an ELISA microplate reader (FluoStar Omega, BMG Labtech) with Omega and Omega MARS software for analysis. A B1-8^hi^ Fab (for NP) or C144 (for SARS2-RBD) antibody served as a normalizer control, and its dilutions were included on each plate. EC₅₀ values were calculated using a four-parameter nonlinear regression model (GraphPad Prism v10.1.1) with the settings: [agonist] vs. response-variable slope (four parameters); bottom = 0; Hillslope > 0; and top equal to the experiment-specific upper plateau of the normalizer control antibody or plasma sample, which was defined as saturation for at least three consecutive dilution steps. The curve fit was constrained to an upper limit corresponding to the maximal optical density achieved by the normalizer control to limit inter-plate and interexperiment variability.

### Biolayer interferometry (BLI) analysis for epitope-mapping

BLI assays were performed as described previously.^8^ Briefly, we used the Octet Red instrument (ForteBio) at 30 °C with shaking at 1,000 r.p.m. For epitope mapping assays, the Protein A biosensor (ForteBio, 18-5010) was used according to the manufacturer’s protocol for a classical sandwich assay, as detailed below: (1) Sensor Check: Sensors were immersed in buffer (ForteBio buffer, 18-1105) for 30 seconds. (2) Capture of First Antibody: Sensors were immersed in buffer with 10 μg/mL first antibody for 10 minutes. (3) Baseline: Sensors were immersed in buffer alone for 30 seconds. (4) Blocking: Sensors were immersed in buffer with 10 μg/mL IgG isotype control (3BNC117) for 5 minutes. (5) Baseline: Sensors were again immersed in buffer for 30 seconds. (6) Antigen Association: Sensors were immersed in buffer with 20 μg/mL RBD for 5 minutes. (7) Baseline: Sensors were immersed in buffer alone for 30 seconds. (8) Association with Second Antibody: Sensors were immersed in buffer for 5 minutes with second antibody at 10 μg/mL. Curve fitting was performed using the ForteBio Octet data analysis software.

## QUANTIFICATION AND STATISTICAL ANALYSIS

Statistical details, including the number of mice or cells per group (n), number of replicates, mean values (center bars), and statistical significance levels, are provided in the figure legends. No specific method was used to assess whether the data met the assumptions of the statistical tests. Statistical significance was determined using GraphPad Prism 10. Data were considered statistically significant at *P ≤ 0.05, **P ≤ 0.01, ***P ≤ 0.001 and ****P ≤ 0.0001.

