## Supplemental Figures for "B lymphocytes that enter the germinal center late preferentially differentiate into memory cells that recognize subdominant epitopes"

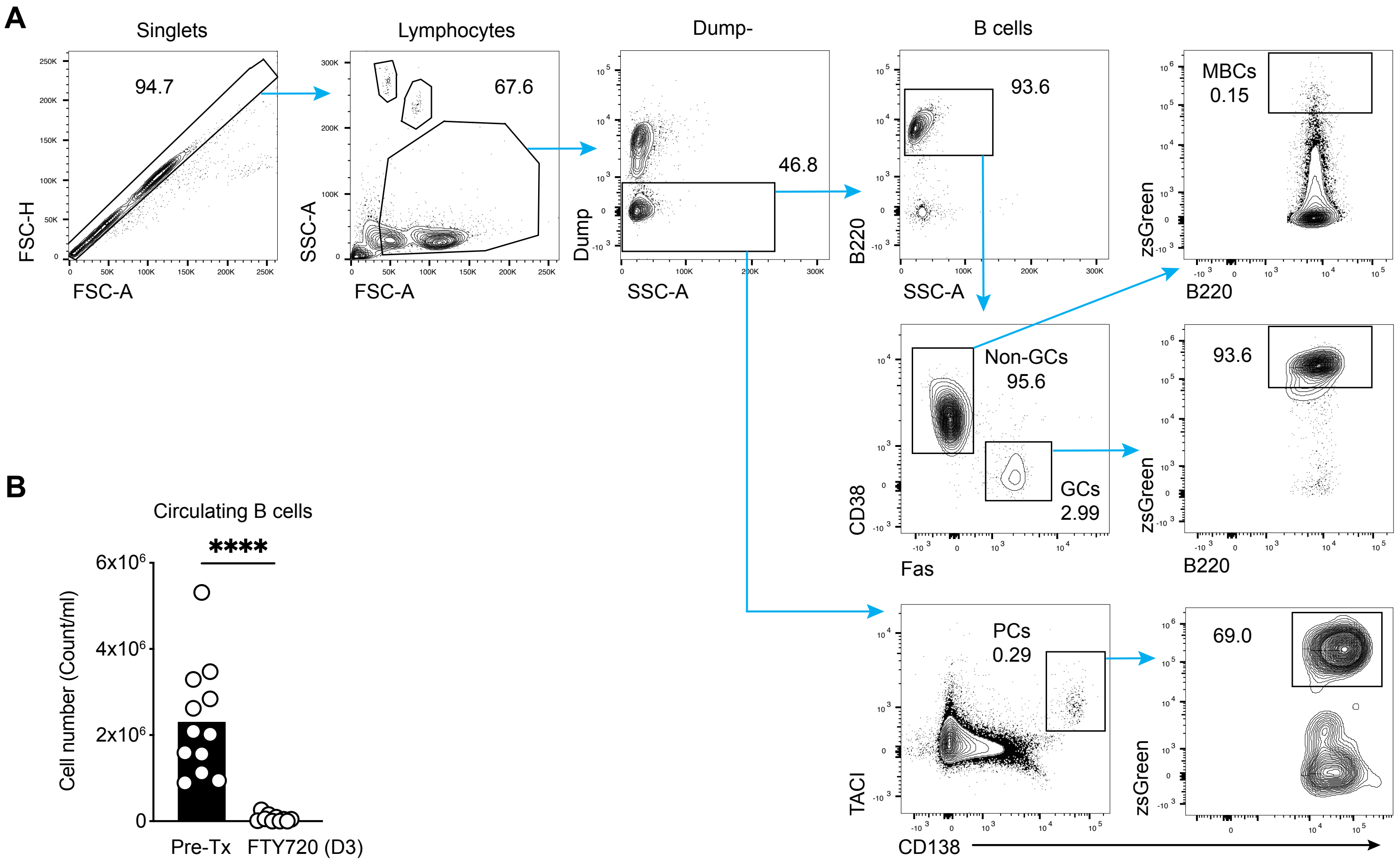
A
Singlets
94.7
FSC-H
FSC-A
Lymphocytes
67.6
SSC-A
FSC-A
Dump-
46.8
Dump
SSC-A
B cells
93.6
B220
SSC-A
MBCs
0.15
zsGreen
B220
Non-GCs
95.6
GCs
2.99
CD38
Fas
93.6
zsGreen
B220
PCs
0.29
TACI
CD138
69.0
zsGreen
B
Circulating B cells
****
6x10⁶
4x10⁶
2x10⁶
0
Cell number (Count/ml)
Pre-Tx
FTY720 (D3)

**Figure S1**

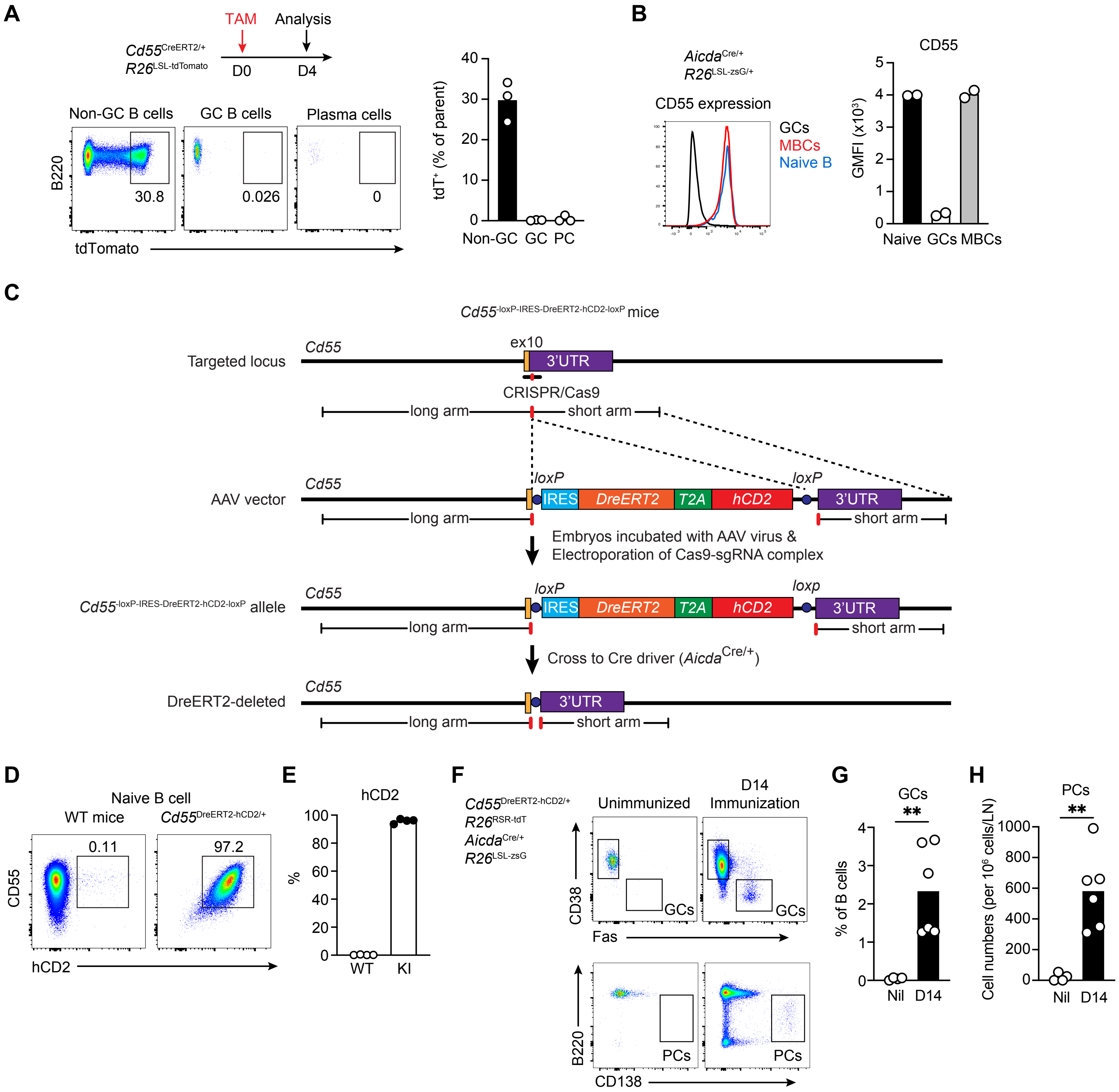
A
TAM
Analysis
Cd55CreERT2/+
R26LSL-tdTomato
D0
D4
Non-GC B cells
GC B cells
Plasma cells
B220
30.8
0.026
0
tdTomato
tdT+ (% of parent)
Non-GC
GC
PC
B
AicdaCre/+
R26LSL-zsG/+
CD55 expression
GCs
MBCs
Naive B
CD55
GMFI (x10³)
Naive
GCs
MBCs
C
Cd55-loxP-IRES-DreERT2-hCD2-loxP mice
Targeted locus
Cd55
ex10
3'UTR
CRISPR/Cas9
long arm
short arm
AAV vector
loxP
IRES
DreERT2
T2A
hCD2
Embryos incubated with AAV virus &
Electroporation of Cas9-sgRNA complex
Cd55-loxP-IRES-DreERT2-hCD2-loxP allele
loxp
Cross to Cre driver (AicdaCre/+)
DreERT2-deleted
D
Naive B cell
WT mice
Cd55DreERT2-hCD2/+
CD55
0.11
97.2
hCD2
E
hCD2
%
WT
KI
F
Cd55DreERT2-hCD2/+
R26RSR-tdT
AicdaCre/+
R26LSL-zsG
Unimmunized
D14
Immunization
CD38
Fas
GCs
B220
CD138
PCs
G
GCs
% of B cells
Nil
D14
H
PCs
Cell numbers (per 10⁶ cells/LN)
Nil
D14

**Figure S2**

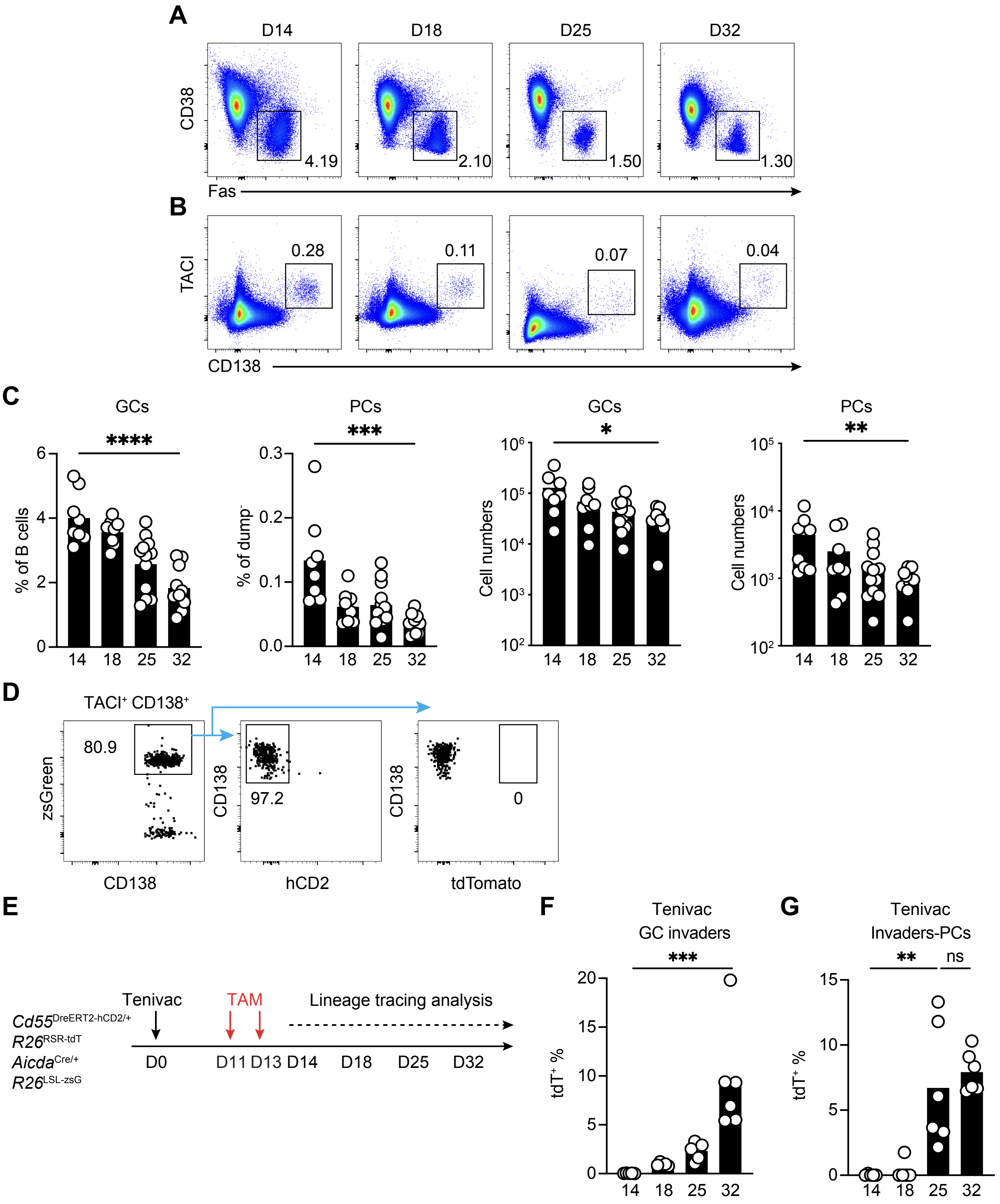
A
D14
D18
D25
D32
CD38
Fas
4.19
2.10
1.50
1.30
B
TACI
CD138
0.28
0.11
0.07
0.04
C
GCs
PCs
% of B cells
% of dump⁻
Cell numbers
****
***
*
**
14 18 25 32
D
TACI⁺ CD138⁺
zsGreen
CD138
80.9
hCD2
97.2
tdTomato
0
E
Tenivac
TAM
Lineage tracing analysis
Cd55DreERT2-hCD2/+
R26RSR-tdT
AicdaCre/+
R26LSL-zsG
D0
D11 D13 D14
D18
D25
D32
F
Tenivac
GC invaders
tdT⁺ %
G
Tenivac
Invaders-PCs
ns

**Figure S3**

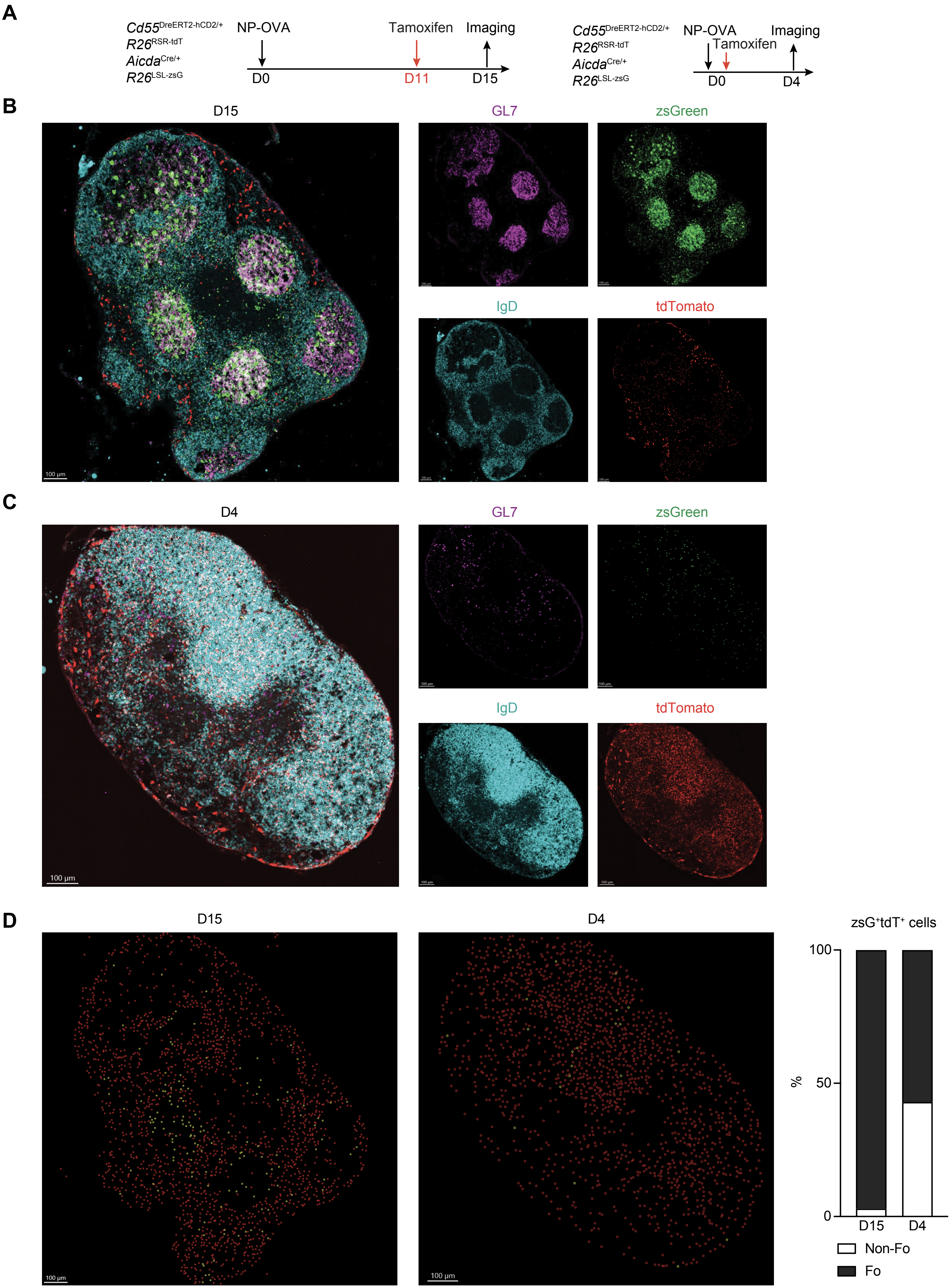
A
Cd55DreERT2-hCD2/+
R26RSR-tdT
AicdaCre/+
R26LSL-zsG
NP-OVA
Tamoxifen
Imaging
D0
D11
D15
Cd55DreERT2-hCD2/+
R26RSR-tdT
AicdaCre/+
R26LSL-zsG
NP-OVA
Tamoxifen
Imaging
D0
D4
B
D15
GL7
zsGreen
IgD
tdTomato
100 µm
C
D4
GL7
zsGreen
IgD
tdTomato
100 µm
D
D15
D4
100 µm
zsG+tdT+ cells
%
100
50
0
D15
D4
Non-Fo
Fo

**Figure S4**

**A**

$$1 - \frac{\dfrac{\text{invaders \%}}{\text{Naive B cell labelling\% / 100}} - \text{invaders \%}}{\text{founders \%}} = \text{Purity of founders}$$

**B**

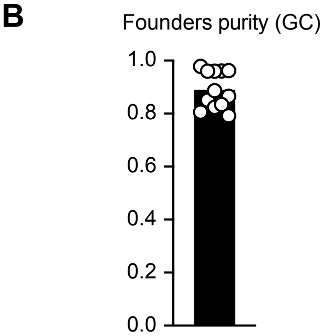


**C**

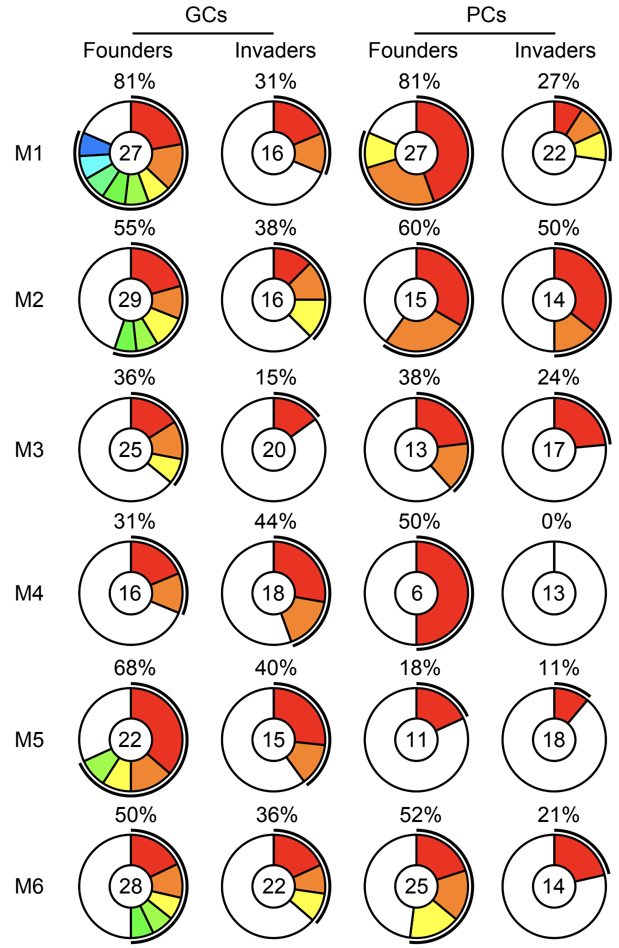


**D**

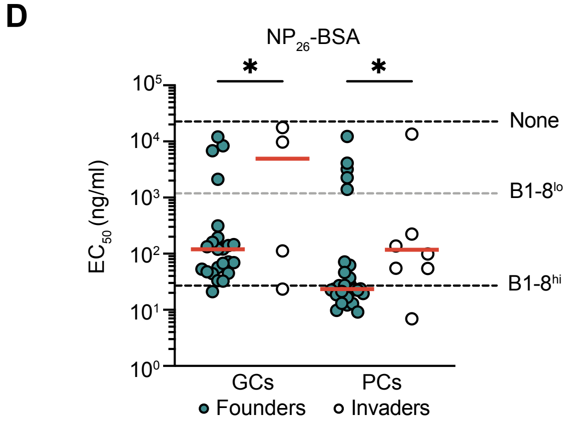


**E**

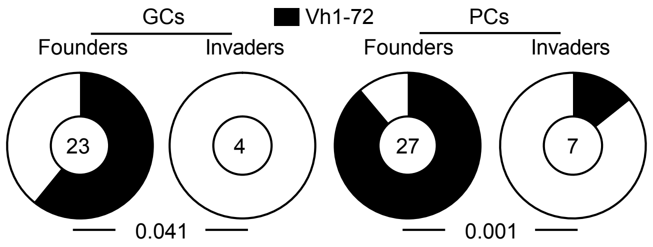


**Figure S5**

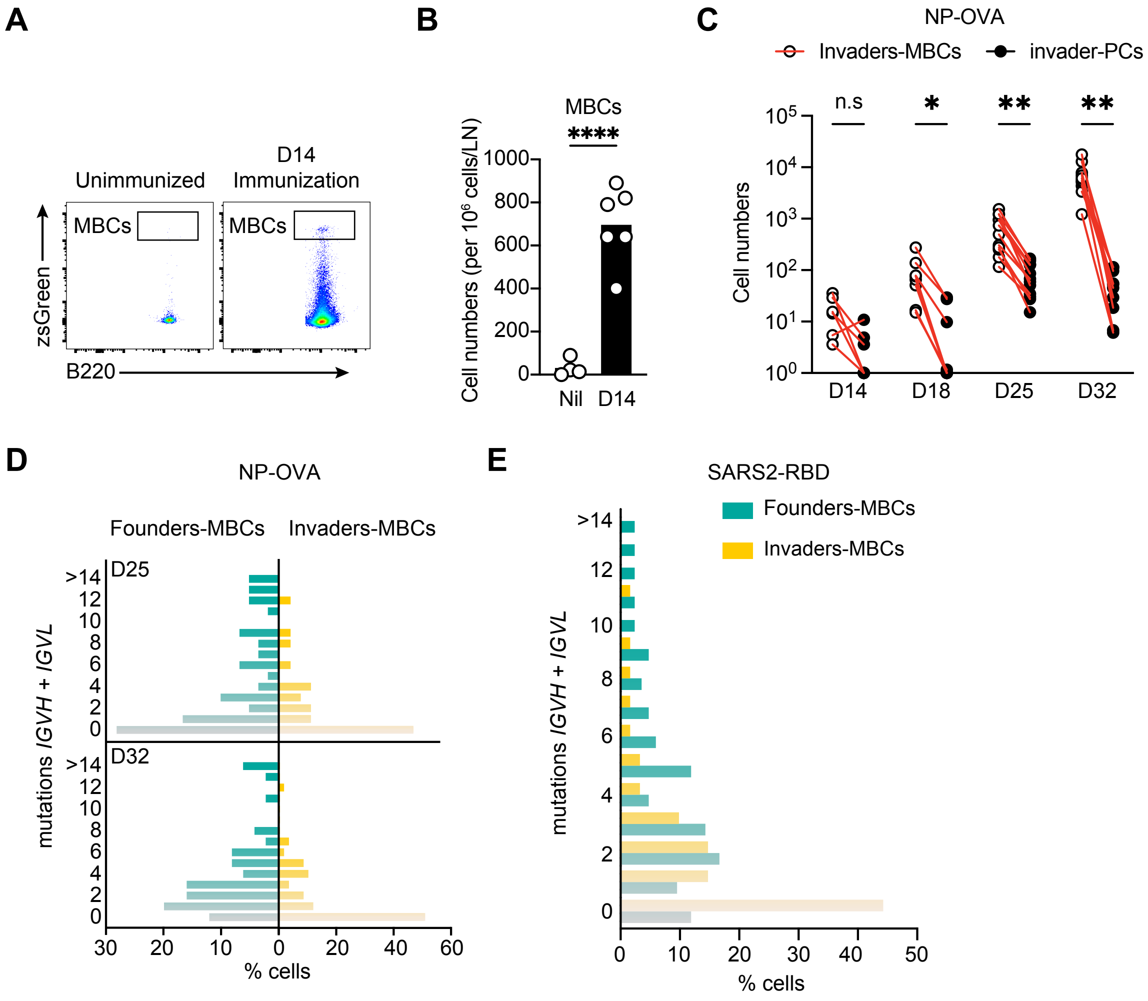
A
Unimmunized
D14
Immunization
MBCs
MBCs
zsGreen
B220
B
MBCs
****
Cell numbers (per 10^6 cells/LN)
1000
800
600
400
200
0
Nil
D14
C
NP-OVA
Invaders-MBCs
invader-PCs
n.s
*
**
**
Cell numbers
D14
D18
D25
D32
D
NP-OVA
Founders-MBCs
Invaders-MBCs
D25
D32
mutations IGVH + IGVL
% cells
E
SARS2-RBD
Founders-MBCs
Invaders-MBCs
mutations IGVH + IGVL
% cells

### Figure S6

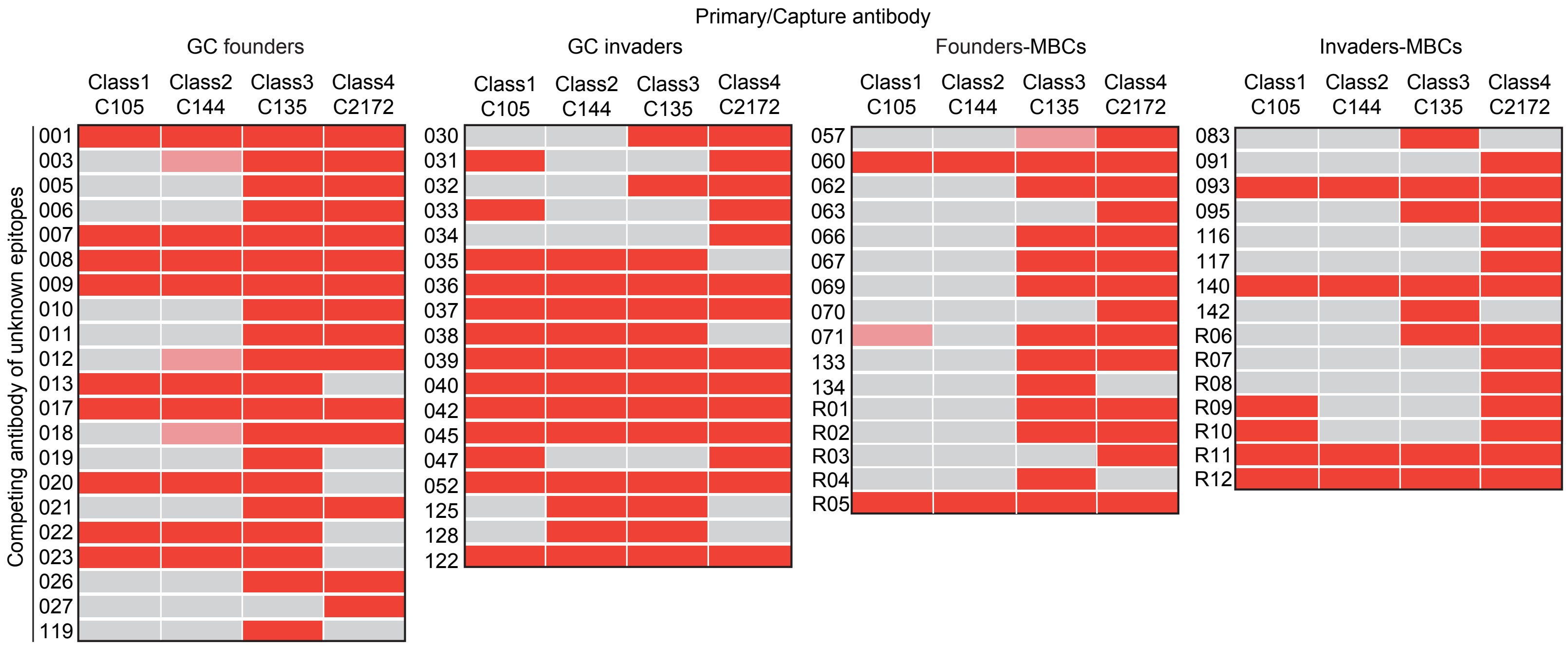
Primary/Capture antibody
GC founders
GC invaders
Founders-MBCs
Invaders-MBCs
Class1 C105
Class2 C144
Class3 C135
Class4 C2172
Competing antibody of unknown epitopes
001 003 005 006 007 008 009 010 011 012 013 017 018 019 020 021 022 023 026 027 119
030 031 032 033 034 035 036 037 038 039 040 042 045 047 052 125 128 122
057 060 062 063 066 067 069 070 071 133 134 R01 R02 R03 R04 R05
083 091 093 095 116 117 140 142 R06 R07 R08 R09 R10 R11 R12

**Figure S7**

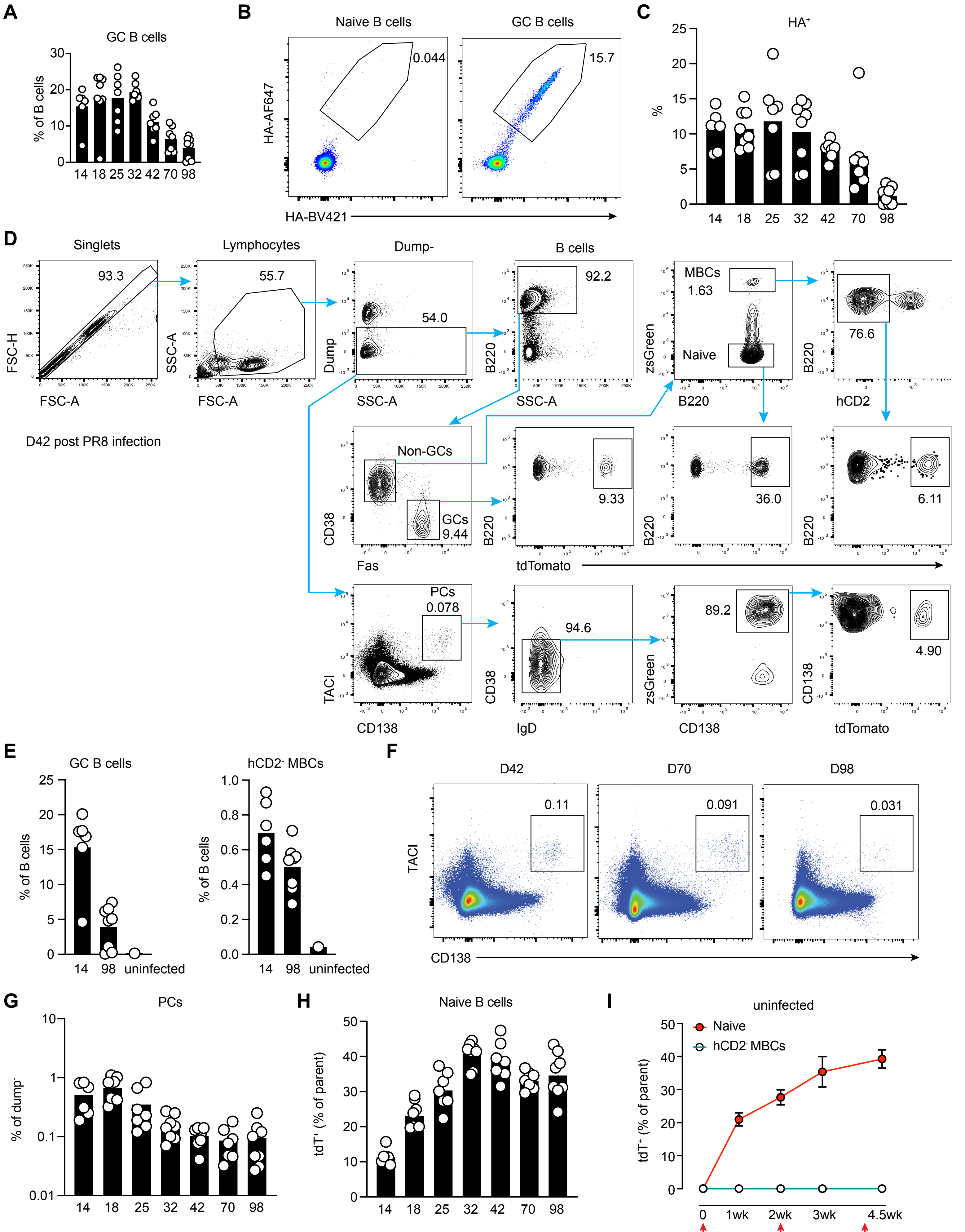

A
GC B cells
% of B cells
14 18 25 32 42 70 98
B
Naive B cells
GC B cells
HA-AF647
HA-BV421
0.044
15.7
C
HA+
%
14 18 25 32 42 70 98
D
Singlets
Lymphocytes
Dump-
B cells
93.3
55.7
54.0
92.2
MBCs
1.63
Naive
76.6
FSC-H
FSC-A
SSC-A
Dump
B220
zsGreen
hCD2
D42 post PR8 infection
Non-GCs
GCs
9.44
9.33
36.0
6.11
CD38
Fas
tdTomato
PCs
0.078
94.6
89.2
4.90
TACI
CD138
IgD
E
GC B cells
hCD2- MBCs
% of B cells
14 98 uninfected
F
D42
D70
D98
0.11
0.091
0.031
G
PCs
% of dump-
H
Naive B cells
tdT+ (% of parent)
I
uninfected
Naive
hCD2- MBCs
0 1wk 2wk 3wk 4.5wk
TAM every 2 weeks


**Figure S8**

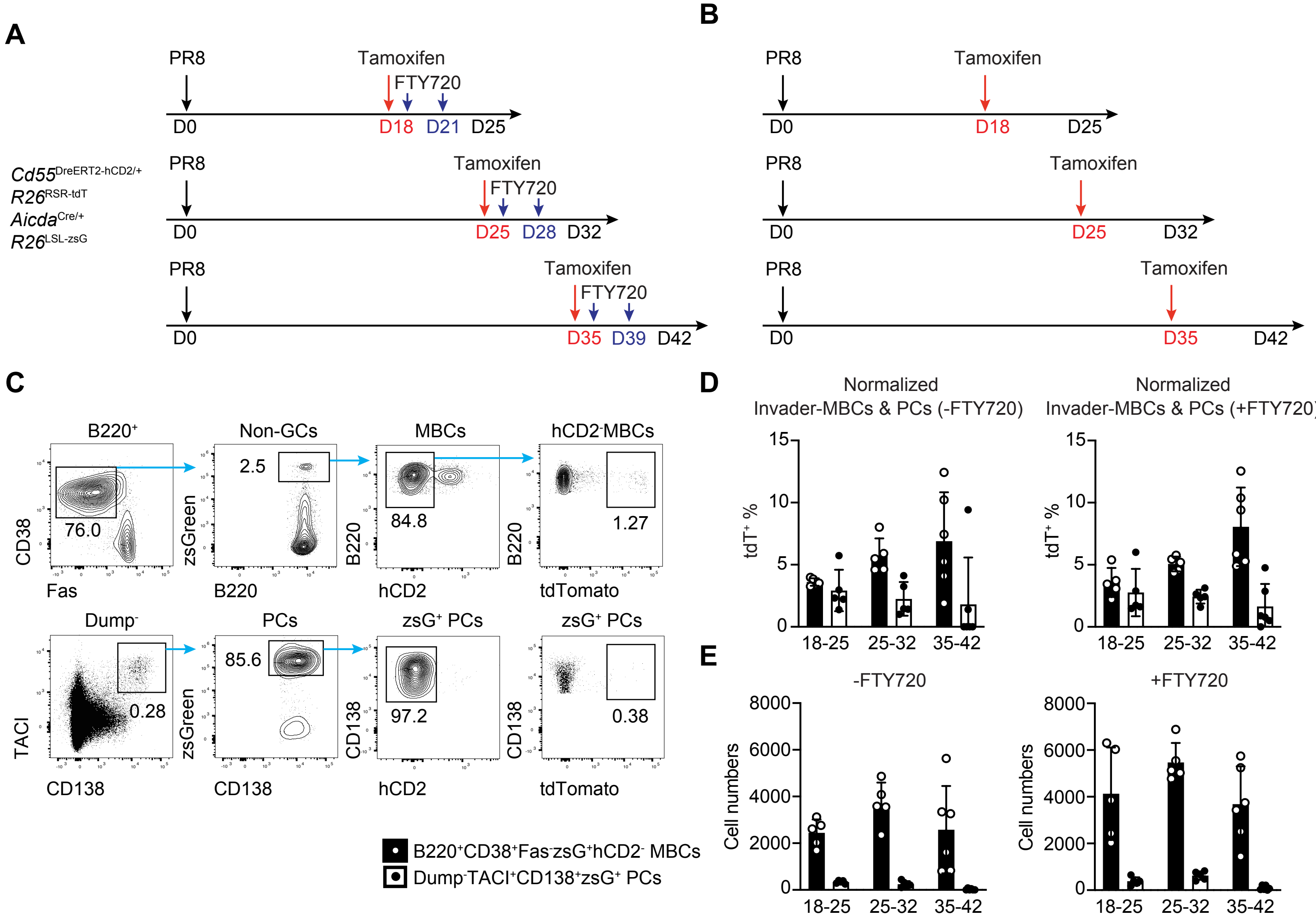
A
B
C
D
E
PR8
Tamoxifen
FTY720
D0
D18
D21
D25
D28
D32
D35
D39
D42
Cd55DreERT2-hCD2/+
R26RSR-tdT
AicdaCre/+
R26LSL-zsG
B220+
Non-GCs
MBCs
hCD2- MBCs
CD38
Fas
76.0
zsGreen
B220
2.5
hCD2
84.8
tdTomato
1.27
Dump-
PCs
zsG+ PCs
TACI
CD138
0.28
85.6
97.2
0.38
B220+CD38+Fas-zsG+hCD2- MBCs
Dump-TACI+CD138+zsG+ PCs
Normalized
Invader-MBCs & PCs (-FTY720)
Invader-MBCs & PCs (+FTY720)
tdT+ %
15
10
5
0
18-25
25-32
35-42
-FTY720
+FTY720
Cell numbers
8000
6000
4000
2000

**Figure S9**
